# An improved auxin-inducible degron system preserves native protein levels and enables rapid and specific protein depletion

**DOI:** 10.1101/585927

**Authors:** Kizhakke Mattada Sathyan, Brian D. McKenna, Warren D. Anderson, Fabiana M. Duarte, Leighton Core, Michael J. Guertin

**Affiliations:** Biochemistry and Molecular Genetics Department, University of Virginia, Charlottesville, Virginia, United States of America; Center for Public Health Genomics, University of Virginia, Charlottesville, Virginia, United States of America; Cancer Center, University of Virginia, Charlottesville, Virginia, United States of America; Department of Stem Cell and Regenerative Biology, Harvard University, Cambridge, Massachusetts, United States of America; Department of Molecular and Cell Biology, University of Connecticut, Storrs, Connecticut, United States of America

**Keywords:** Auxin Inducible Degron, Auxin Response Factor, ZNF143, Transcription Factors, RNA Polymerase Pausing

## Abstract

Rapid perturbation of protein function permits the ability to define primary molecular responses while avoiding down-stream cumulative effects of protein dysregulation. The auxin-inducible degron (AID) system was developed as a tool to achieve rapid and inducible protein degradation in non-plant systems. However, tagging proteins at their endogenous loci results in chronic, auxin-independent degradation by the proteasome. To correct this deficiency, we expressed the Auxin Response Transcription Factor (ARF) in an improved inducible degron system. ARF is absent from previously engineered AID systems, but ARF is a critical component of native auxin signaling. In plants, ARF directly interacts with AID in the absence of auxin and we found that expression of the ARF Phox and Bem1 (PB1) domain suppresses constitutive degradation of AID-tagged proteins. Moreover, the rate of auxin-induced AID degradation is substantially faster in the ARF-AID system. To test the ARF-AID system in a quantitative and sensitive manner, we measured genome-wide changes in nascent transcription after rapidly depleting the ZNF143 transcription factor. Transciptional profiling indicates that ZNF143 activates transcription in *cis* and ZNF143 regulates promoter-proximal paused RNA Polymerase density. Rapidly inducible degradation systems that preserve the target protein’s native expression levels and patterns will revolutionize the study of biological systems by enabling specific and temporally defined protein dysregulation.

## Introduction

The function of proteins can be studied in cells using RNAi depletion, loss-of-function mutants, temperature sensitive mutations, small molecule inhibitors, CRISPR interference, or nucleic acid aptamers. The two greatest limitations of these methods are 1) the chronic nature of permanently disrupting function and 2) the limited availability and specificity of inhibitor molecules. Inducible degradation methods, such as auxin-inducible degron (AID) (Nishimura et al. 2009) or degradation tag (dTAG) systems (Nabet et al. 2018), can overcome these limitations. Rapidly regulated systems and inhibitors permit measurements of primary changes in molecular, cellular, and organismal phenotypes and subsequent tracking of the cascade that accompanies immediate protein dysregulation. Gene editing techniques now permit endogenous tagging of genes, which should preserve the target gene’s expression levels and patterns. Endogenous fusion of target genes with rapidly inducible degradation tags have the potential to revolutionize the way we study biological systems by providing specific and temporally defined perturbation techniques for any protein of interest.

Auxin is a plant hormone that regulates various aspects of plant growth and development. In plants, auxin signaling triggers a rapid switch between transcriptional repression and transcriptional activation (Calderon-Villalobos et al. 2010; Lavy and Estelle 2016; Li et al. 2016) (Figure S1). There are three key components in this signal transduction system: 1) transport inhibitor response 1 (TIR1), a subunit of a ubiquitin ligase complex that binds to the target substrate; 2) the auxin response transcription factors (ARF), which directly regulate gene expression; and 3) auxin/indole-3-acetic acid (Aux/IAA) proteins, which are destabilized in the presence of auxin-mediated ubiquitination. In the absence of auxin, the domains III&IV of Aux/IAA family of proteins form heterodimeric complexes with the Phox and Bem1 (PB1) domain of ARF proteins (Kim et al. 1997; Ulmasov et al. 1999). Domain I of Aux/IAA interacts with a plant-specific transcriptional co-repressor, Topless (TPL) (Szemenyei et al. 2008). The repressive function of TPL dominates relative to the activation function of ARF transcription factors (Tiwari et al. 2004). In the presence of auxin, TIR1 interacts with domain II of Aux/IAA to facilitate ubiquitination and degradation of Aux/IAA (Dharmasiri et al. 2003; Gray et al. 2001), which liberates ARFs to regulate transcription. This auxin-inducible degron system was engineered to function outside plant cells (Nishimura et al. 2009).

Although the AID system has been widely adopted to degrade tagged proteins of interest, recent studies report auxin-independent depletion of endogenously tagged proteins (Morawska and Ulrich 2013; Nishimura and Fukagawa 2017; Zasadzinśka et al. 2018). For instance, tagging the centromeric histone chaperone, HJURP, results in depletion of more than 90% of protein in the absence of auxin in human cell lines (Zasadzinśka et al. 2018). Chronic depletion has also been reported in chicken cells (Nishimura and Fukagawa 2017) and in yeast cells (Morawska and Ulrich 2013), suggesting that auxin-independent depletion may be a systemic problem when tagging endogenous genes. The extent of factor-dependent depletion is often impossible to evaluate because many studies do not directly compare protein levels in the progenitor and tagged cell lines (Cao et al. 2018; Hoffmann et al. 2016; McKinley et al. 2015). Neither defining a minimal degradation domain of AID nor employing an inducible TIR1 system have overcome this deficiency (Mendoza-Ochoa et al. 2018; Morawska and Ulrich 2013; Samejima et al. 2014). In yeast, auxin-independent depletion of the AID-tagged factors is dependent on cellular TIR1 concentration, suggesting that depletion is due to ubiquitin-mediated degradation (Mendoza-Ochoa et al. 2018).

Herein, we show that AID-tagging of endogenous genes commonly results in auxin-independent and chronic depletion by proteasome-mediated protein degradation. We found that co-expressing the PB1 domain of ARF improved the robustness of the AID system by rescuing auxin-independent degradation and increasing the rate of auxin-induced degradation. Our control experiments also revealed that auxin treatment alone results in the activation of the Aryl Hydrocarbon Receptor (AHR) transcription factor and regulation of AHR target genes. Excluding AHR-responsive genes from downstream differential expression analyses is critical when investigating the activity of AID-tagged transcriptional regulators. Collectively, these improvements enhance the robustness, sensitivity, and specificity of the AID system. We applied ARF-rescue, AID-mediated, rapid degron depletion to the transcription factor ZNF143 to identify the primary ZNF143 response genes and to define a functional role of ZNF143 in transcriptional regulation.

## Results

### Endogenous auxin-inducible degron tagging results in chronic target protein depletion

In non-plant systems, the auxin-mediated degradation system requires the presence of exogenously expressed TIR1 (Nishimura et al. 2009). We generated an HEK293T cell line with the TIR1 gene stably integrated into the AAVS1 locus (Mali et al. 2013; Natsume et al. 2016) (Figure S2). We independently tagged all copies of either ZNF143, TEAD4, and p53 in these progenitor HEK293T-TIR1 cells (Figure 1A). In the absence of auxin, tagging the factors resulted in depletion of the proteins to levels that range from between <3%-15% of endogenous levels, as measured by quantitative Western blots (Figure 1B,C,&D). To determine whether chronic depletion is unique to HEK293T cells, we generated an MCF-7 cell line with TIR1 incorporated heterozygously into the AAVS1 locus and tagged all copies of ZNF143, which also resulted in auxin-independent ZNF143 depletion (Figure 1E). Next, we characterized a previously published AID-tagged CENP-I colorectal epithelial DLD-1 cell line (McKinley et al. 2015). CENP-I is also modestly depleted (Figure 1F). The modest degree of CENP-I depletion may be because CENP-I is an essential protein and a minimal abundance of CENP-I is necessary for cell line viability. Basal depletion of these factors in three distinct cell lines confirms the generality of auxin-independent depletion of endogenously tagged proteins.

**Fig. 1.**
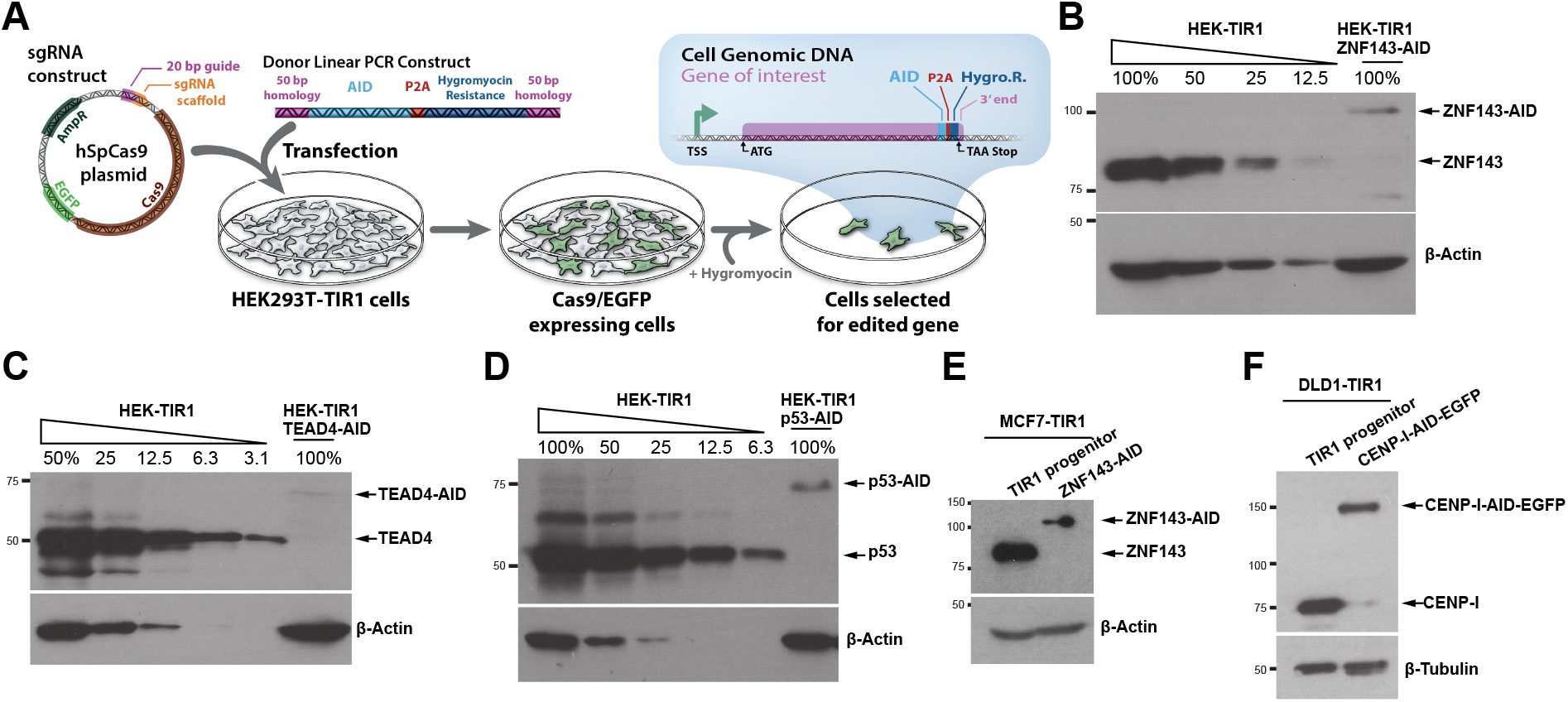
Endogenous tagging of genes with AID results in auxin-independent protein depletion. A) Genes of interest were targeted by cotransfecting sgRNAs (Table 2) targeting the 3*′* end of the coding sequence with a homology-directed repair donor construct. The donor construct contained the AID domain fused to a hygromycin resistance marker; an intervening porcine teschovirus-1 2A (P2A) site liberates the hygromycin resistance marker during protein translation. This construct was flanked by 50 base pair homology regions that correspond to the sequences flanking the sgRNA recognition sequence and Cas9 cleavage site. B,C,&D) We quantified AID-tagged protein abundance with quantitative Western blots and a dilution of the progenitor cell line for the standard curve. Endogenous homozygous tagging of ZNF143 (B), TEAD4 (C), and p53 (D) in HEK293T cells results in auxin-independent chronic protein depletion. MCF-7 (E) and DLD-1 (F) cell lines also exhibit depletion of endogenously tagged ZNF143 (E) and CENP-I (F).

### Proteasome mediated degradation drives auxin-independent depletion of proteins

We next sought to investigate the mechanism by which AID-tagged proteins are depleted by focusing on AID-tagged ZNF143. We performed nascent RNA sequencing (Core et al. 2008; Kwak et al. 2013; Mahat et al. 2016) in the progenitor HEK293T-TIR1 cells and the ZNF143-AID cells. Nascent ZNF143 transcript levels remain unchanged (1.08-fold increase, FDR = 0.27) in the AID-tagged ZNF143 cell line (Figure 2A). Because the auxin-inducible system is dependent upon the proteasome and AID can interact with TIR1 in the absence of auxin (Dharmasiri et al. 2005; Gray et al. 2001; Kepinski and Leyser 2005; Tan et al. 2007), we hypothesized that low protein abundance may be mediated by an auxin-independent interaction with TIR1, ubiquitination, and subsequent proteasome-mediated degradation. We treated ZNF143-AID cells with the proteasome inhibitor MG132 and observed a modest increase in ZNF143 levels after 4.5 hours (Figure S3A). TIR1 depletion also results in higher ZNF143-AID and TEAD4-AID levels (Figure S3B&C). These results indicate that chronic, auxin-independent depletion of AID-tagged proteins is TIR1-mediated and due to proteasomal degradation (Figure S3D).

**Fig. 2.**
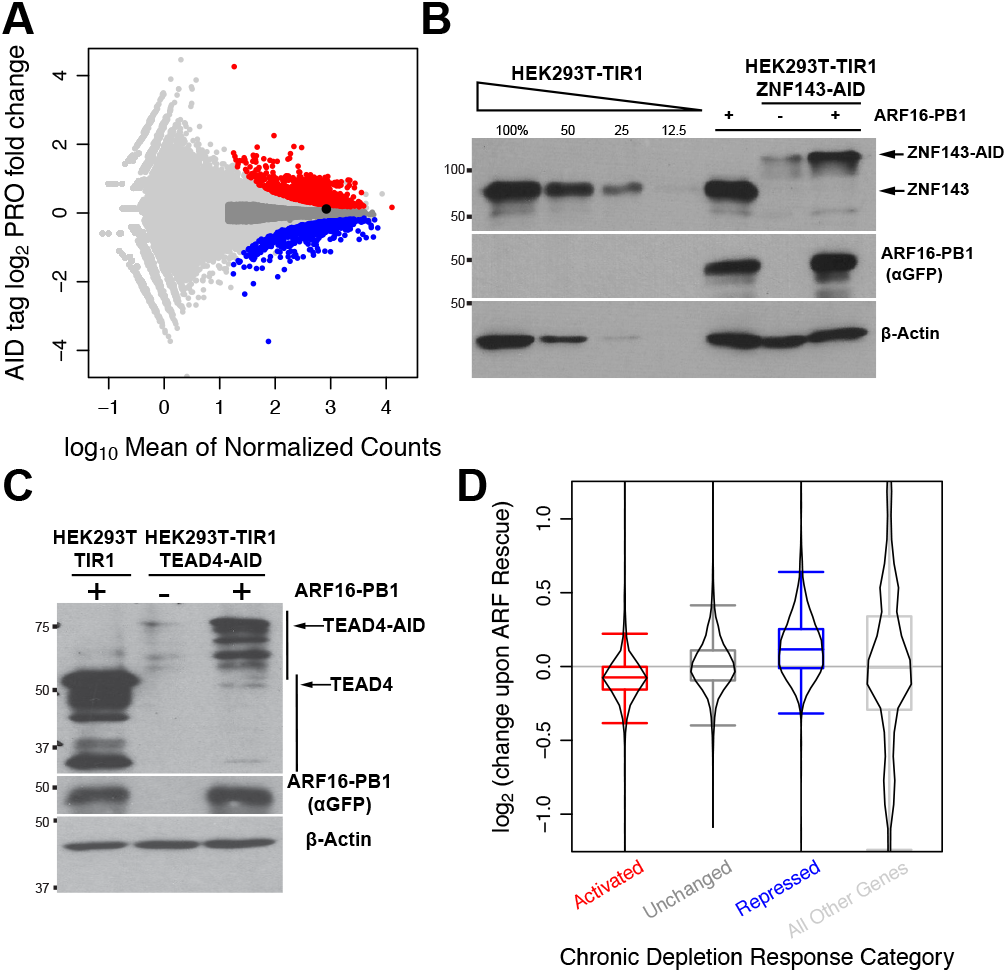
Expressing the PB1 domain of ARF rescues the proteasome-dependent chronic depletion of AID-tagged proteins. A) Red genes are activated upon chronic ZNF143 depletion, blue genes are repressed, dark grey genes are confidently unchanged and expressed at comparable levels to changed genes, and light grey points represent all other genes. ZNF143 (black point) expression is unaffected. B) Stable expression of ARF-PB1 domain in HEK293T-TIR1 cells did not change the protein levels of ZNF143 (lane 5), but increased the stability of ZNF143-AID (lane 7). C) TEAD4 levels also increase upon ARF16 expression. D) Genes that are activated upon chronic ZNF143 depletion in panel A are, on average, repressed upon ARF rescue and genes that are repressed upon chronic depletion are activated upon ARF rescue.

### Auxin Response Factor rescues auxin-independent proteasomal degradation

TIR1 and AID proteins can directly interact in the absence of auxin *in vitro* (Dharmasiri et al. 2005; Kepinski and Leyser 2005; Tan et al. 2007). However, degradation is tightly and robustly regulated in plants (Chapman and Estelle 2009) (Figure S1). We proposed that expression of an ARF protein, which is an AID-interaction partner, may confer protection of the tagged protein from auxin-independent ubiquitination and degradation. To test this hypothesis, we supplemented the engineered AID system by expressing the AID-interacting PB1 domain of ARF. We reconstituted the system by expressing the PB1 domain of *Oryza sativa* ARF16 and ARF25, based on yeast two-hybrid experiments that quantified interaction of these ARFs with IAA proteins (Shen et al. 2010). Moreover, ARF16 and ARF25 harbor conserved charged residues, corresponding to K944, D994, and D998 of ARF16 (Wang et al. 2007), at critical positions within the ARF/IAA binding interface (Korasick et al. 2014; Nanao et al. 2014). Transfection of either eGFP-ARF25-MR-PB1 or eGFP-ARF16-PB1 stabilizes TEAD4-AID, with ARF16 promoting a higher degree of TEAD4 stability (Figure S4A&B). These results prompted us to generate HEK293T-TIR1-ZNF143-AID cells with stable genetic integration and expression of ARF16-PB1. This strategy restored ZNF143 levels to over 50% of untagged levels (Figure 2B and Figure S4C&D). Similarly, we found that genetic integration and expression of eGFP-ARF16-PB1 stabilized endogenously tagged TEAD4-AID (Figure 2C). In contrast, stable expression of eGFP-ARF16-PB1 did not alter ZNF143, TEAD4, or p53 protein levels in HEK293T or HEK293T-TIR1 cells (Figure S4E).

Transcriptional output is a quantitative measure of ARF-mediated functional rescue of ZNF143. We performed nascent RNA transcriptional profiling (Core et al. 2008) with the three successive genetically modified HEK293T cells: progenitor TIR1 cells, chronic ZNF143-depleted AID-tagged cells, and ARF-rescued ZNF143-AID cells. Chronic ZNF143 depletion resulted in activation of 1188 genes and repression of 774 genes at a false discovery rate (FDR) of 0.001 (Figure 2A). Next, we analyzed the raw changes in expression upon ARF rescue to determine whether rescuing ZNF143 stability can functionally rescue gene expression profiles. Of the 1188 activated genes upon chronic ZNF143 depletion, 899 (76%) decrease their expression upon ARF rescue (Figure 2D). Of the 774 chronically repressed genes, 561 (72%) increase their expression upon ARF rescue (Figure 2D). These changes are consistent with a functional rescue of gene expression upon ARF rescue of ZNF143 stability.

### Auxin Response Factor rescue mediates rapid aux-in-inducible degradation

We treated ZNF143-AID and TEAD4-AID cells with 500 *μ*M auxin to determine how the ARF16 rescue affects inducible depletion. Auxin treatment induces degradation of both ZNF143-AID and TEAD4-AID in a time dependent manner (Figure 3A&B). Importantly, the rate of degradation of ZNF143 was increased upon ARF16-PB1 rescue when compared to cells not rescued with ARF16-PB1 (Figure 3C&D). To test whether ARF16-PB1 rescue affects the synthesis rate of ZNF143-AID, and thus contributing to the perceived degradation rate, we simultaneously blocked new protein synthesis with cycloheximide at the time of auxin treatment (Figure S5). Upon blocking protein synthesis, the ZNF143-AID protein degraded faster in the presence of ARF16-PB1 (Figure S5). Therefore, ARF expression promotes faster degradation kinetics and ARF does not influence protein synthesis rate.

**Fig. 3.**
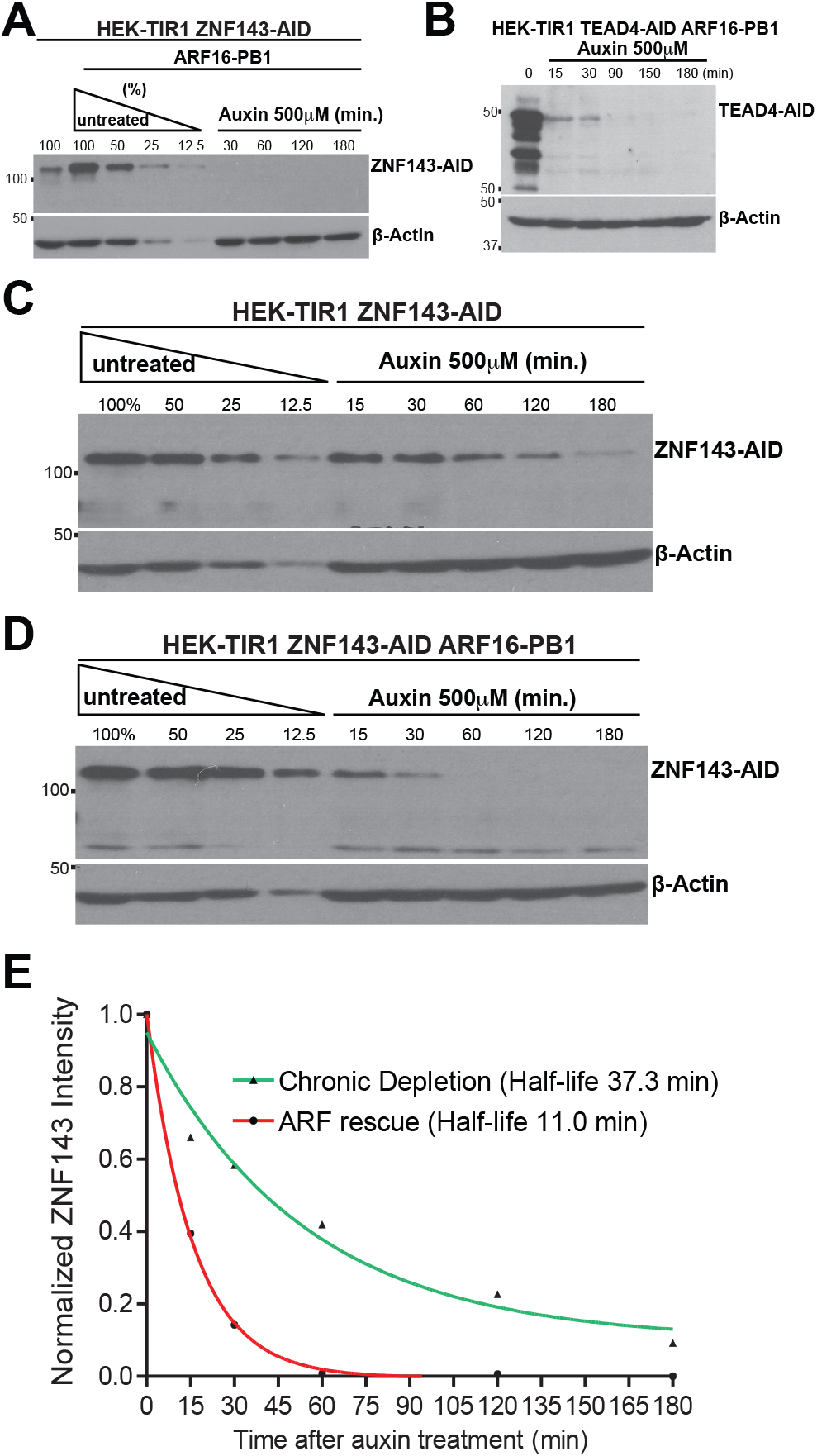
ARF rescue promotes faster degradation kinetics upon addition of auxin. ZNF143 (A) and TEAD4 (B) are rapidly depleted upon auxin treatment. C) A quantitative Western blot measures the time dependent degradation of ZNF143-AID in the presence of 500*μ*M auxin. Note that ZNF143 is chronically depleted in these cells and a longer exposure was needed to quantify auxin-induced reduction of ZNF143-AID levels. D) A quantitative Western blot measures 500*μ*M auxin-induced ZNF143-AID degradation after ARF16-PB1 rescue. E) We measured the intensity of ZNF143-AID bands from panels C&D using densitometry and fit the data using nonlinear regression and a one-phase decay equation.

### Auxin Response Factor interacts with AID to rescue AID tag stability in mammalian cells

To test the specificity of the ARF16-PB1 rescue, we mutated ARF16 residues within the interaction domain interface that are critical for its interaction with IAA17 (Korasick et al. 2014; Nanao et al. 2014) (Figure 4A&B). We converted K944, D994, and D998 to Alanine in the ARF16-PB1-MT. In *Arabidopsis thaliana*, the corresponding mutations abolish ARF homodimerization and heterodimerization with IAA17 (Korasick et al. 2014). We tested whether this mutant is capable of rescuing AID tagged protein stability in HEK293T-TIR1 cells. Chronic depletion of AID-tagged ZNF143 and TEAD4 was rescued with wild-type ARF16-PB1, but not with ARF16-PB1-MT (Figure 4C&D). However, we note that the ARF16-PB1 mutant protein is not as abundant as the wild type ARF16-PB1 (Figure 4C&D), presumably because dimerization can stabilize exogenous ARF16-PB1. This result suggests that rescue of stability is mediated by the interaction between ARF16-PB1 and AID. To confirm a physical interaction, we performed a co-immunoprecipitation experiment. We found that NLS-mCherry-AID and eGFP-ARF16-PB1 interact in the absence of TIR1 (Figure 4E). However, the ARF16 mutations are sufficient to disrupt the ARF16/AID interaction (Figure 4E). The reciprocal IP confirmed the AID/ARF16-PB1 interaction and we were unable to detect an interaction of TIR1 with AID (Figure 4F). These data indicate that an ARF16-PB1/AID interaction mediates AID stability in the improved ARF-AID degron system.

**Fig. 4.**
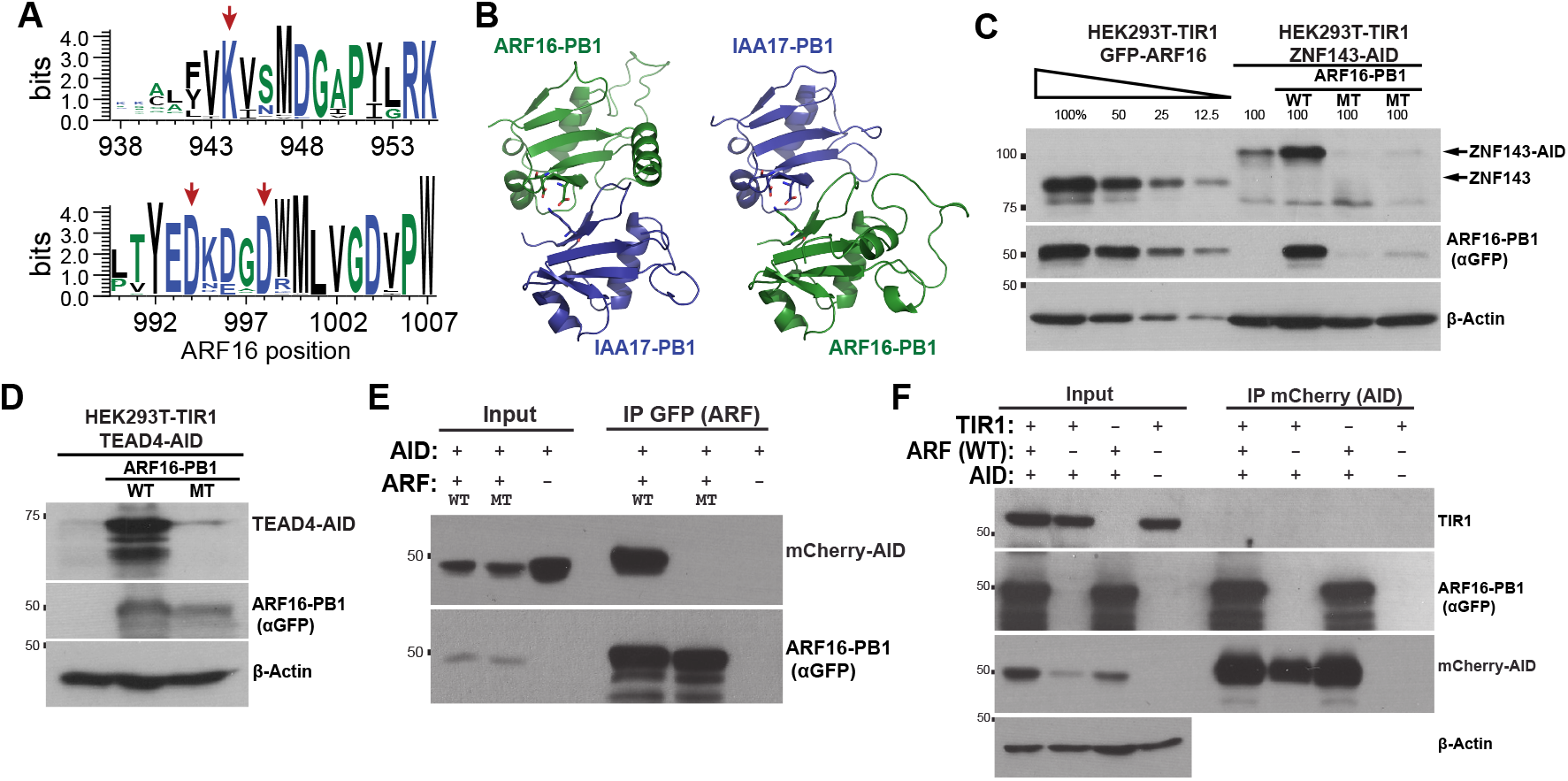
The interaction of ARF-PB1 and AID mediates rescue of endogenous protein levels. A) Positions indicated by the red arrows are highly conserved within the AUXIAA family (Pfam Family: PF02309). A seed alignment of the Aux-IAA family was generated using Pfam (El-Gebali et al. 2018) and the resultant FASTA files were visualized using WebLogo (Crooks et al. 2004). Positions on the x-axis are relative to ARF16 residue numbers. B) The amino acids D994 and D998 of ARF16 and K114 of IAA17 are shown in the left panel. The amino acid side chains corresponding to D183 and D187 of IAA17 and K944 if ARF16 are highlighted in the right panel. Note that IAA17/ARF16 heterodimeric complexes are shown in both orientations and that these domains of IAA17 and ARF16 are within the same protein domain family. IAA17 and ARF16 sequences were modeled (Waterhouse et al. 2018) into each chain of an *Arabidopsis thaliana* ARF5 homodimeric structure (PDB entry: 4CHK (Nanao et al. 2014)). These mutations in ARF16, which disrupt the electrostatic binding interface, fail to rescue chronic ZNF143 (C) and TEAD4 (D) degradation. E) These mutations disrupt this co-immunoprocipitation of mCherry-AID with eGFP-tagged ARF16-PB1. Consistent with lower stability of the ARF16-MT in panels C&D, the mutant GFP-ARF16-PB1 plasmid was transfected at a concentration three times higher than the wild type to achieve comparable expression of each protein. F) ARF16-PB1 is detected upon mCherry-AID immunoprecipitation, however we were unable to detect TIR1 associating with AID.

### Expression of ARF-PB1 and TIR1 prior to AID-tagging preserves native protein levels

Rescuing chronic ZNF143 depletion does not result in full recovery of protein levels and approximately 25% of the genes that are differentially expressed in the chronic depletion remain dysregulated upon ARF-rescue (Figure 2D). To further improve the system and preserve native levels of the tagged protein, we constitutively expressed ARF16-PB1 prior to AID tagging. We generated a bicistronic construct with ARF-PB1 and TIR1 and an intervening P2A site, which separates the two proteins during translation. Genetic incorporation of this construct into the AAVS1 locus resulted in two independent clones that express ARF16-PB1 and TIR1 (Figure 5A). ZNF143 AID-tagging of both ARF-PB1/TIR1 HEK293T progenitor cell lines results in ZNF143 protein levels that are comparable to the parental lines (Figure 5B&C).

**Fig. 5.**
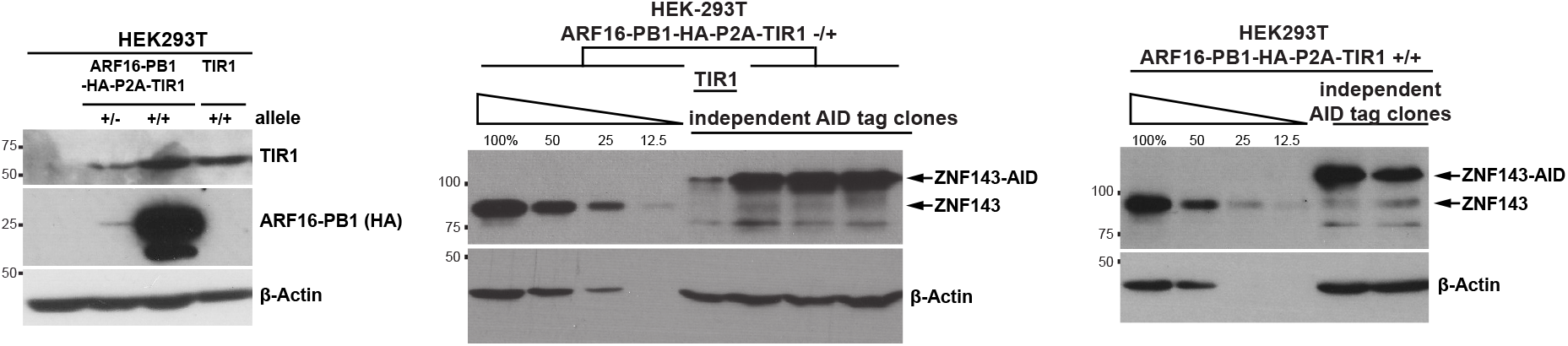
Expression of ARF-PB1 prior to AID-tagging preserves ZNF143 protein levels. A) We generated two progenitor cell lines that express HA-tagged ARF16-PB1 and TIR1 from the same promoter, separated by a P2A site. Unmodified HEK293T cells are within lane 1. The construct is incorporated into at least one allele of AAVS1 (lane 2) and all AAVS1 alleles (lane 3). Note that HEK293T cells are not strictly diploid. Lane 4 is the original TIR1-expressing progenitor cell line. B) Three independent ZNF143-AID clones (lanes 6-8) have comparable protein levels as the heterozygous ARF-PB1/TIR1 progenitor cells. Consistent with previous figures, TIR1 expression alone compromises ZNF143-AID protein levels. C) Two independent ZNF143-AID clones (lanes 5&6) derived from the homozygous ARF-PB1/TIR1 cells preserve ZNF143 protein levels.

### ZNF143 activates transcription in *cis*

We directly and quantitatively assessed auxin-induced perturbation of ZNF143 by measuring genome-wide transcriptional changes upon auxin treatment. Conventional transcriptional profiling, such as RNA-seq, require mature RNA to accumulate or degrade to detect changes in transcription. Delayed detection poses a challenge when measuring the immediate transcriptional response upon rapid protein depletion. To overcome this limitation, we measured nascent RNA using PRO-seq (Core et al. 2008; Kwak et al. 2013; Mahat et al. 2016). PRO-seq quantifies the immediate effect that degradation has upon transcribing RNA Polymerases. ZNF143 was first characterized as a sequence-specific activator as measured by reporter assays (Schuster et al. 1995). In order to identify direct ZNF143 target genes and primary response genes, we performed genome-wide nascent RNA profiling after 90 minutes of auxin treatment in the chronically depleted and ARF-rescued ZNF143-AID cell lines. We define *primary effect genes* as immediately regulated upon rapid ZNF143 depletion and *direct gene targets* as primary response genes that ZNF143 regulates in *cis*. Many genes are activated and repressed upon auxin treatment in both backgrounds (Figure 6A). ZNF143 binding, as measured by ChIP-seq (Consortium et al. 2012), is enriched proximal to the repressed gene class in the ARF rescue background (Kolmogorov–Smirnov two-sided p-value = 1.1e^*−*16^) and not the activated gene class (p-value = 0.047) (Figure 6B). In the chronic ZNF143 depletion background, the auxin-repressed gene class is not significantly closer to ZNF143 binding sites (p-value = 0.022) and genes within the activated gene class tend to be further away from ZNF143 binding sites (p-value = 0.0017) (Figure 6B). This supports recent genomic data, which found that many transcription factors (ER, GR, PPAR*γ*, NF*κ*B, HSF) specialize to exclusively activate or repress transcription in *cis* (Carroll et al. 2006; Duarte et al. 2016; Guertin et al. 2014; Reddy et al. 2009; Schmidt et al. 2016, 2015; Step et al. 2014; Vockley et al. 2016).

**Fig. 6.**
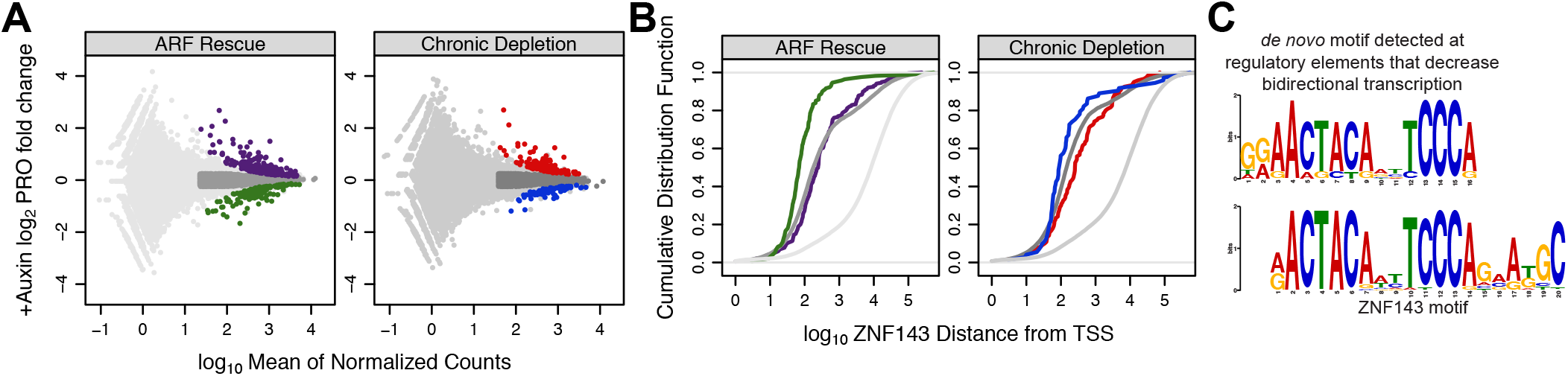
ZNF143 is a canonical transcriptional activator. A) Genes are activated and repressed upon auxin treatment in both the chronic ZNF143-depleted background and the ARF-rescue of ZNF143 degradation background. Purple points are genes that increase expression and green genes decrease expression upon auxin treatment in the ARF-rescued background. Dark grey genes are expressed at comparable levels as activated and repressed genes, but they are confidently unchanged when auxin is added. B) Cumulative distribution plots quantify the relationship between the closest ZNF143 ChIP-seq peak and the start sites of genes within the regulated classes from panel A; trace colors correspond to panel A categories. ZNF143 binding sites are closer to auxin-repressed genes only in ARF-rescue background, suggesting that ZNF143 functions to activate transcription in *cis*. C) The ZNF143 motif was found *de novo* exclusively at regulatory elements that decrease bidirectional transcription upon auxin treatment.

Transcription factors can direct bidirectional transcription at their binding sites (Hah et al. 2013; Kim et al. 2010). Bidirectional transcription is a feature of gene promoters and enhancers (Core et al. 2014) and a measure of regulatory element activity. We tested whether we could detect changes in bidirectional nascent RNA production upon ZNF143 depletion. We implemented a discriminative regulatory-element detection method (dREG) (Wang et al. 2019) to identify promoters and enhancers *de novo*. We performed differential bidirectional transcription analysis to identify regulatory elements that increase and decrease transcriptional activity upon auxin treatment (Figure S6). Next, we performed *de novo* motif analysis (Bailey et al. 2009) within the regulatory elements that increase or decrease bidirectional transcription. The canonical ZNF143 motif was exclusively found in the auxin-repressed regulatory elements (Figure 6C). Taken together with the integrative ChIP-seq/PRO-seq analysis from Figure 6A&B, we conclude that ZNF143 activates transcription of proximal genes and enhancers. These results serve as direct evidence that we are not only depleting ZNF143 protein levels, but we are functionally perturbing ZNF143 activity using the ARF-AID system.

### ZNF143 targets are more responsive to perturbation upon ARF16 rescue

Of the 168 genes that are classified as *repressed* upon auxin treatment in the ARF-rescue background, 167 genes have a net negative change in auxin-induced gene expression in the chronic depletion background (Figure S7A). Upon auxin treatment, eighty-seven percent (146/168) of the genes have a greater magnitude of response in the ARF-rescue background compared to chronic ZNF143-depleted cells (Figure S7A). These data show that the ARF-rescue is more sensitive to detect auxin-induced changes in ZNF143-dependent transcription compared to the chronically depleted background. Seventy-eight percent (57/73) of the auxin-repressed genes in the chronic ZNF143-depletion background are categorized as *repressed* in the rescue as well (Figure S7B); seventy percent (40/57) are repressed to a greater magnitude in the rescue (Figure S7B). This analysis indicates that expressing ZNF143 at near-endogenous levels is necessary to detect a robust transcriptional response upon ZNF143 depletion.

### Auxin treatment activates the Aryl Hydrocarbon Receptor response

In order to determine whether this system could be generally applied to study transcription factor function, we performed a control experiment to test whether auxin treatment alone affects transcription of human genes. Few genes are repressed upon auxin treatment (Figure S8A), regardless of FDR thresholds. However, over a range of FDR thresholds we consistently observe that the activated genes (Figure S8) are enriched in Aryl Hydrocarbon Receptor (AHR) binding sequences in their promoters and the most enriched pathway for this gene set is the Aryl Hydrocarbon Receptor pathway (q-value = 0.002) (Kuleshov et al. 2016). We find that AHR binding (Lo and Matthews 2012) is enriched proximal to the activated gene class (p-value = 3.6e^*−*11^) and not the repressed gene class (p-value = 0.74) (Figure S8B&C). This control experiment highlights the importance of filtering aryl hydrocarbon receptor response genes from analyses when using any AID system to study transcriptional response.

### ZNF143 regulates paused RNA Polymerase density

Transcription can be regulated at various steps (Fuda et al. 2009; Scholes et al. 2017), including chromatin opening (Adelman et al. 2006; Guertin and Lis 2010; Morris et al. 2014), pre-initiation complex formation/stability and RNA Polymerase II (Pol II) recruitment (Stargell and Struhl 1996), Pol II initiation (Esnault et al. 2008; Govind et al. 2005; Sakurai and Fukasawa 2000), Pol II pausing and release (Adelman and Lis 2012; Marshall and Price 1992), and elongation (Ardehali et al. 2009). General transcription machinery and cofactors directly catalyze these steps, but these factors are targeted to DNA by sequence-specific transcription factors, such as ZNF143. We sought to determine which transcription step(s) ZNF143 targets by characterizing the change in RNA Pol II profiles after rapid ZNF143 depletion (Figure S9). We found that the repressed gene class, which represents direct ZNF143 targets, show dramatic changes in the pause region compared to the gene body (Figures 7A&B, S10, and S11).

**Fig. 7.**
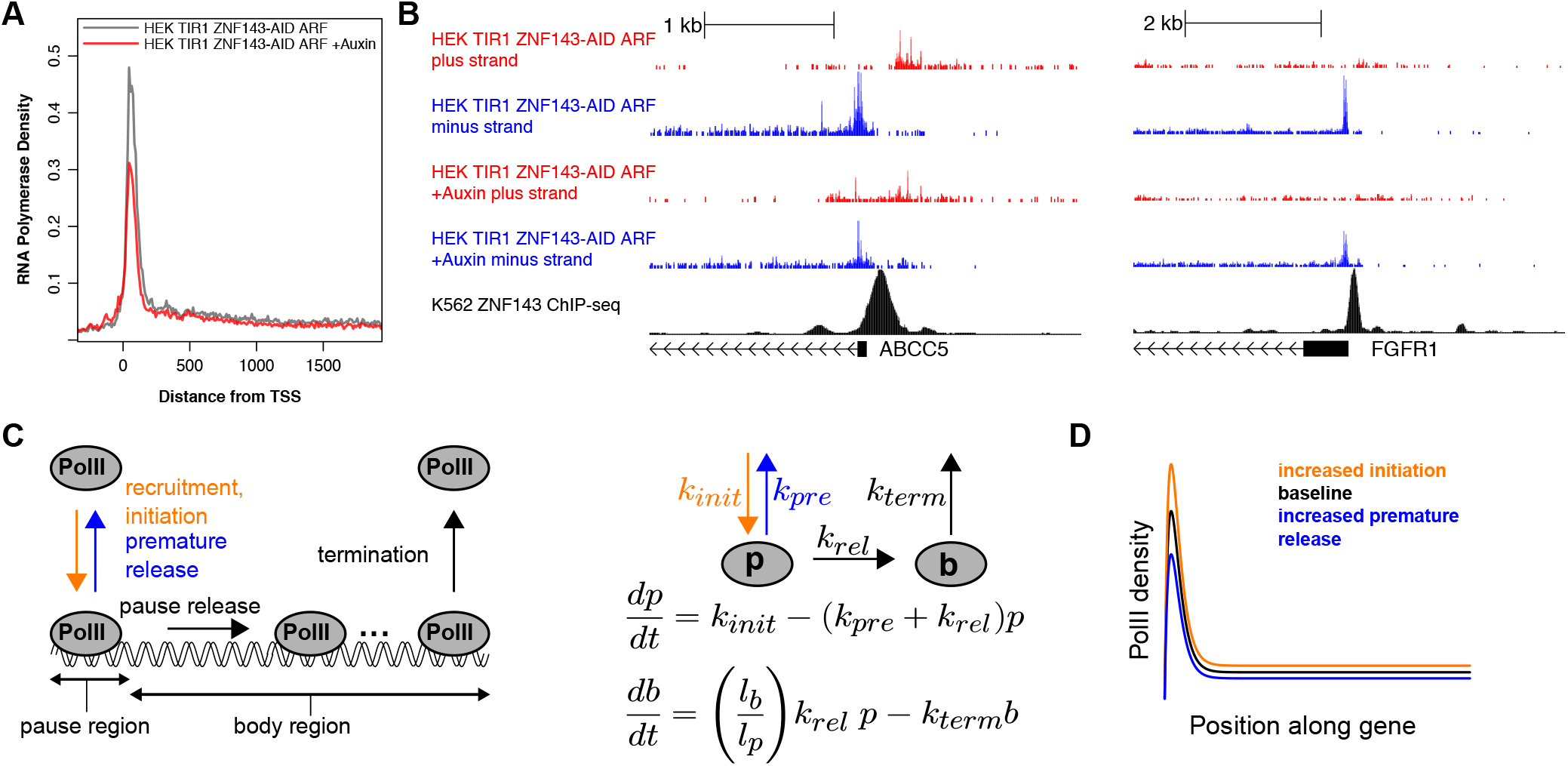
ZNF143 regulates initiation/recruitment or pause stability. A) The composite profile of Pol II density at all auxin-repressed genes indicates that Pol II pause density is compromised upon ZNF143 depletion. B) Individual genes (ABCC5 and FGFR1) show compromised Pol II density in the pause region. C) Model structure and key variables are highlighted in this schematic. A mathematical formulation of the two component model in which *p* refers to Pol II density at the pause region and *b* refers to the density at the gene body region. D) This plot represents a steady state simulation for a reference model (black), a model in which transcriptional initiation was increased by 25% (orange), and a model in which premature pause release was increased by 60% (blue). The peak of the smooth curve was set to the steady state pause level and the plateau of the curve was set to the steady state gene body level. Note that this plot captures the preferential effect on the pause region as compared to the gene body region.

We implemented a mathematical modeling approach to better understand the potential mechanisms underlying decreased pause/body densities for repressed genes following ZNF143 inhibition. We formulated a two compartment model with dynamics for a pause region and gene body region. Model parameters include rate constants for transcriptional initiation/Pol II recruitment, premature pause release, pause release into productive transcription elongation, and termination of transcription (Figure 7C). We implemented parameter sensitivity analyses to determine if any model parameters could account for our experimental results. Our analysis showed that only changes in initiation and premature pause release could account for a large magnitude of Pol II density change in the pause region relative to the gene body. Increased initiation resulted in increases for both the pause region and the gene body region. In contrast, increases in premature release resulted in decreases for both the pause region and the gene body region (Figure 7D). These results suggest that decreases in the pause and gene body regions observed following ZNF143 inhibition could be accounted for by either decreases in initiation or increases in non-productive pause release. We used the model to further investigate the conditions under which the change in the pause region could be greater than the change in the gene body region for a variation in either the initiation or premature release rate. Our analysis showed that such preferential changes in the pause region will always be observed when the ratio of Pol II density in the pause region relative to the to the gene body, known as the pause index, is greater than a value of one. Moreover, the relative pause change was predicted by the model to be identical to the pause index, which is consistent with our result that the pause index at repressed genes is only modestly changed at genes that are repressed upon ZNF143 depletion (Figure S12). These combined experimental and modeling results suggest that ZNF143 regulates pausing density by facilitating RNA Polymerase II initiation or preventing premature dissociation of paused RNA Polymerase II.

## Discussion

Expression of ARF-PB1 improves the auxin inducible degradation system. First, ARF-PB1 interacts with the AID-tagged protein to prevent degradation in the absence of auxin, thus the tagged protein’s abundance is more representative of native levels. Secondly, the rate of auxin-induced depletion is increased in the ARF-AID system. We demonstrated the power of the ARF-AID system by rapidly depleting the transcription factor ZNF143 and quantifying genome-wide changes RNA Polymerase density.

### Advantages of rapidly inducible degron systems

Protein function can be studied by rapidly inducible degron systems in cases where translational fusion of the degron tag does not disrupt protein function or protein stability. These systems provide advantages in interpretation of protein function because measurements taken immediately after protein dysregulation can be attributed directly to the protein of interest. In contrast, other techniques that are general, such as RNAi and genetic knockout, do not provide opportunities to assay phenotypes immediately after protein depletion due to the gradual or chronic nature of dysregulation. The newly developed dTAG system (Nabet et al. 2018) provides comparable advantages as ARF-AID and exogenous expression of two additional proteins is not required. Endogenous tagging of BRD4 with FKBP12^*F*36*V*^ dTAG did not result in dramatic protein depletion in the absence of the inducible degradation molecule, dTAG-13 (Nabet et al. 2018). As the dTAG and ARF-AID systems become more widely adopted, we look forward to studies that systematically compare these different degron technologies.

We found that auxin treatment alone causes undesired transcriptional changes at AHR target genes, but it is unclear whether dTAG-13 treatment results in off-target changes in cellular phenotypes. These types of side effects can be abrogated by including proper control experiments and by depleting target proteins by multiple independent methods.

### Possible mechanisms of ARF-mediated AID stabilization and rapid degradation

ARF transcription factors are a critical component of the plant auxin-response system (Guilfoyle and Hagen 2007). Herein we find that ARF expression is an important component of engineered auxin-inducible degron systems. The ARF-AID system confers two distinct advantages 1) ARF expression limits auxin-independent degradation of target proteins and 2) ARF expression promotes more rapid auxin-inducible degradation of AID-tagged proteins. ARF and Aux/IAA proteins each harbor a conserved PB1 domain that can homodimerize or heterodimerize (Kim et al. 1997; Ulmasov et al. 1999). Mutations that interfere with ARF/AID interaction fail to rescue chronic, auxin-independent AID-degradation. TIR1 binds to domain II of Aux/IAA (Dharmasiri et al. 2003; Gray et al. 2001) and ARF binds to domain III and IV; therefore, ARF and TIR1 do not directly compete for the same binding surface of Aux/IAA. Therefore, an ARF/AID interaction may cause conformational changes within AID that inhibit its interaction with auxin-unbound TIR1 (Figure 8).

**Fig. 8.**
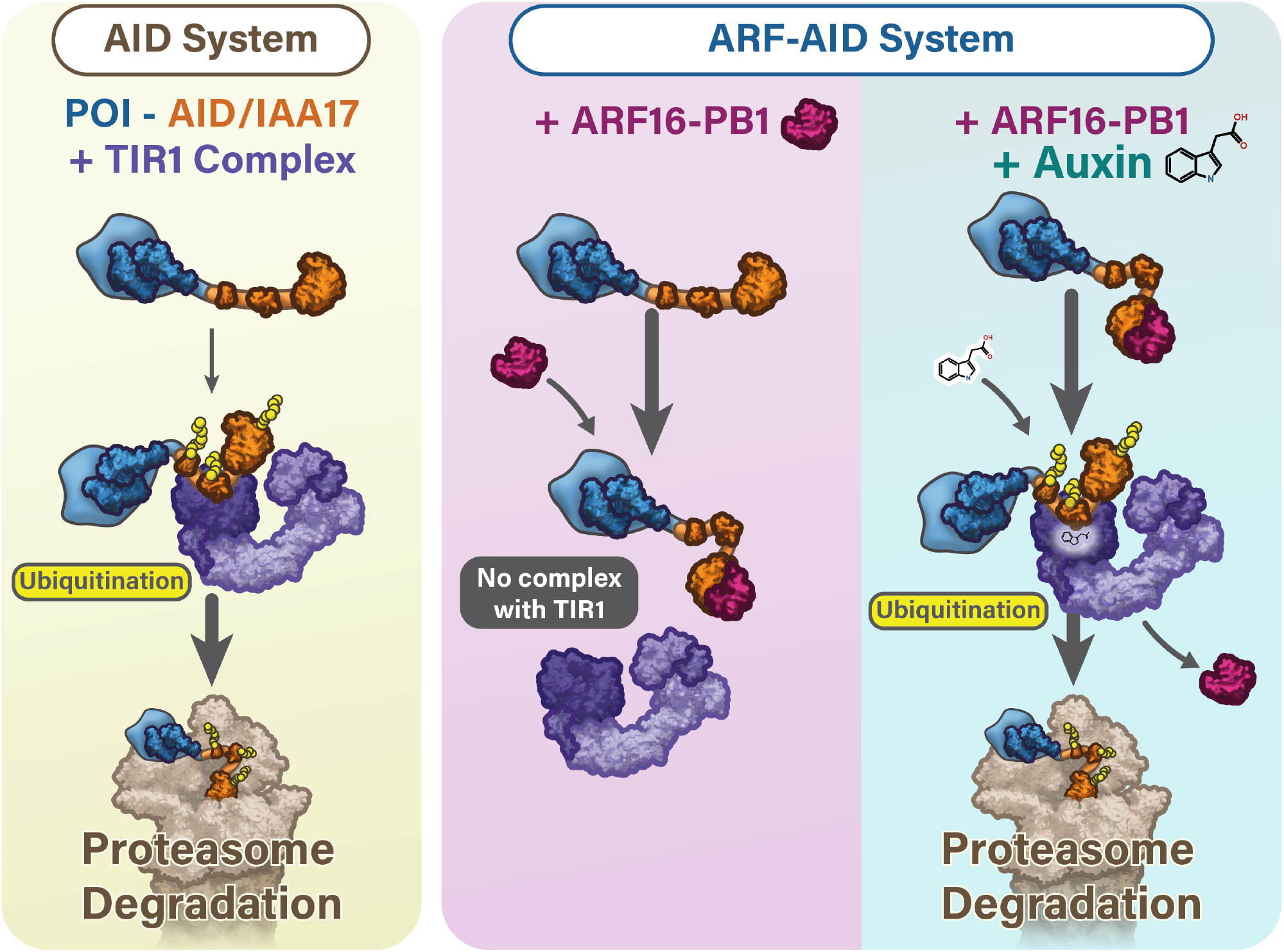
The ARF-AID system preserves protein of interest (POI) expression levels in the absence of auxin and auxin treatment induces rapid degradation. A) The classical AID system can result in auxin-independent degradation of the AID tagged proteins. B) In the ARF-AID system, ARF-PB1 binds to AID to prevent TIR1 association with AID, which prevents auxin-independent AID degradation by the ubiquitin-mediated proteasomal pathway. Auxin facilitates the interaction of TIR1 with AID, promotes dissociation of ARF, and subsequent ubiquitination and proteasome-mediated degradation of the AID-tagged protein.

Auxin-inducible degradation of ZNF143 is more rapid in the ARF-AID system. There are many plausible and non-mutually exclusive possibilities for the observed difference in degradation kinetics. The most simple explanation is that promoting the stability of the AID-tagged protein increases its concentration, which directly affects degradation kinetics. Alternatively, ARF binding to AID may promote more efficient ubiquitination by exposing target residues to the auxin-bound TIR1 complex. ARF binding may cause allosteric changes in domain II of AID that promote a higher affinity interaction with auxin-bound TIR1 compared to ARF-unbound AID. Recent work has shown that different Aux/IAA proteins can interact with TIR1 in the absence of auxin to influence the association of auxin with TIR1 (Villalobos et al. 2012). Although we did not detect an interaction of TIR1 and IAA17 in Figure 4, we cannot dismiss such an interaction in cells. Therefore, the ARF-AID system may promote faster kinetics because ARF binding affects AID structure, which in turn modulates the affinity of TIR1 and auxin. Future studies and systematic comparisons are needed to determine the mechanisms that contribute to a more rapid auxin-inducible response.

### Molecular functions of ZNF143

Despite incredible advances in our understanding of the mechanisms of eukaryotic transcription and developments in systems biology, accurately predicting direct target genes and primary response genes of transcription factors remains a challenge. Proximal binding of a transcription factor to a gene is neither necessary nor sufficient to modulate gene expression. A fundamental question remains: how do transcription factors discriminate between genes in the genome in which to regulate? To begin to address this question we must first define the set of genes regulated by the transcription factor of interest. Technical limitations preclude experimentally identifying a comprehensive set of primary response genes for the vast majority of transcription factors, because we cannot rapidly induce or rapidly repress their activity. Perturbation methodologies that can be universally applied to any gene, such as RNAi, require days to efficiently deplete protein. This time-frame of depletion poses a major barrier to understanding transcription factor function, because secondary (and beyond) effects dominate conventional depletion/knockout methods. Herein, we show that applying the ARF-AID system to study transcription factor function overcomes these challenges.

The *Xenopus laevis* homolog of ZNF143, Staf (selenocysteine tRNA gene transcription activating factor), was first cloned and characterized nearly 25 years ago (Schuster et al. 1995). This original report characterized the binding site of Staf within a regulatory element of the tRNA^*Sec*^ gene and characterized the activator function of Staf using reporter assays (Schuster et al. 1995). Recent transcriptional profiling upon siRNA-mediated ZNF143 depletion identified many activated and repressed genes (Ngondo-Mbongo et al. 2013). ZNF143 depletion caused twice as many genes to decrease expression relative to the number of genes that increased expression. The authors concluded that ZNF143 is primarily an activator, but they note that ZNF143 may be involved in repression. Our results corroborate the activation function of ZNF143 and indicate that although many genes are activated upon immediate ZNF143 depletion, ZNF143 does likely not act in *cis* to mediate repression. Importantly, we inhibited ZNF143 for only 90 minutes and we measured nascent RNA levels, so the repressive role of ZNF143 cannot be attributed to the post-primary response of ZNF143 dysregulation. Alternative mechanisms of immediate indirect repression, such as squelching (Guertin et al. 2014; Schmidt et al. 2016; Step et al. 2014), may be responsible for the observed repressive role of ZNF143. We further characterized ZNF143’s role in activation and found that ZNF143 functions to control paused RNA Pol II density. Mathematical modeling indicates that ZNF143 either positively regulates RNA Polymerase initiation or prevents non-productive dissociation of paused Pol II.

ZNF143 is also involved in chromatin looping of distal enhancers to promoters (Bailey et al. 2015a). Enhancer-promoter looping frequently and preferentially occurs at promoters containing paused RNA Pol II (Ghavi-Helm et al. 2014). Therefore, we propose a model whereby ZNF143 directly regulates the amount of paused Pol II on a given promoter, which facilitates enhancer looping.

Rapidly inducible systems, such as hormone signaling and heat shock response, have contributed greatly to our understanding of transcriptional regulation. The success of these models, in part, is because the regulatory processes can be triggered instantaneously and tracked. New technologies and inhibitors that permit rapid and specific protein dysregulation promise to revolutionize the study of complex regulatory mechanisms.

## Methods

### Cell lines

HEK293T cells were purchased from ATCC and were grown in DMEM with 10% FBS, Penicillin/Streptomycin and 5% glutamine. MCF7 cells were purchased from ATCC and were grown in DMEM with 10% FBS and Penicillin/Streptomycin. CENP-I-AID-eGFP DLD1-OsTIR1 cells were generated in Ian Cheeseman’s lab (McKinley and Cheeseman 2017) and grown in RPMI1640 media with 10% FBS and Penicillin/Streptomycin.

### Plasmids and constructs

OsTIR1 was integrated into AAVS1 locus of the HEK293T cells using the CMV-OsTIR1-PURO plasmid from Masato Kanemaki (pMK232, Addgene #72834) (Natsume et al. 2016). OsTIR1 was integrated into the genome by CRISPR-Cas9 mediated repair using an sgRNA targeting AAVS1 safe harbor locus cloned into pSpCas9(BB)-2A-GFP from Feng Zhang (PX458, Addgene #48138) (Mali et al. 2013; Natsume et al. 2016; Ran et al. 2013). The ARF16-PB1 domain (amino acid 878-1055) and ARF25 (MR and PB1 domain, amino acid 369-889, Os12t0613700-01) genes were codon optimized for humans and synthesized from Bio Basic Inc, New York. We inserted ARF25-MR-PB1 into the eGFP-C2 vector. A nuclear localization signal (NLS) was added at the N-terminus of the ARF16-PB1 domain and NLS-ARF16-PB1 was inserted into eGFP-C2 vector digested with XhoI and HindIII by cold fusion cloning (pMGS36, Addgene #126581). The CMV-OsTIR1-PURO plasmid was digested with AfeI and we inserted the codon optimized ARF16-PB1 domain separated by P2A from Os-TIR1 to generate the ARF16-PB1-HA-P2A-OsTIR1 construct (pMGS46, Addgene #126580). This plasmid was co-transfected with AAVS1 sgRNA (pMGS7, Addgene #126582) and we selected puromycin resistant clones to generate the homozygously integrated transgenic cell line. The resulting transgenic HEK293T-TIR1 and ARF16-PB1-HA-P2A-OsTIR1 cell lines were used to tag transcription factors with AID tag using the CRISPR-Cas9 system. The NLS-mCherry-AID plasmid was constructed by digesting pCDNA5 vector with PmeI enzyme and inserted the NLS-mCherry-AID fragment using cold fusion cloning (System Biosciences).

### Endogenous AID tagging in HEK293T cells

Endogenously AID tagged TEAD4, ZNF143 and p53 cells were generated using CRISPR mediated gene editing. sgRNAs that target the 3*′* end of the respective coding sequences were cloned into hSpCas9 plasmid (PX458, Addgene Plasmid #48138) (Ran et al. 2013). The linear donor was generated by PCR and gel purified from a plasmid harboring a synthetic AID-P2A-Hygromycin insert (pMGS54, Addgene #126583). We amplified the insertion using primers that contain 50 nucleotide homology tails. The primers contained 5*′* phosphorothioate modifications to increase PCR product stability in the cell (Zheng et al. 2014). The primers used for making PCR donor fragments are reported in Table 1. HEK293T cells were co-transfected with 1 *μ*g of CRISPR/Cas9-sgRNA plasmid and 400 ng of linear donor PCR product using Lipofectamine 3000 in a 6 well plate. Cells were expanded into 10 cm plates two days after transfection. The knock-in cells were selected by treating with 200 *μ*g/ml Hygromycin B three days after transfection. Individual clones were selected and confirmed by Western blotting and Sanger sequencing of PCR amplicons. The sgRNAs from Table 2 were used for targeting the 3*′* end (C-terminus of the protein) of the indicated genes.

**Table 1.**
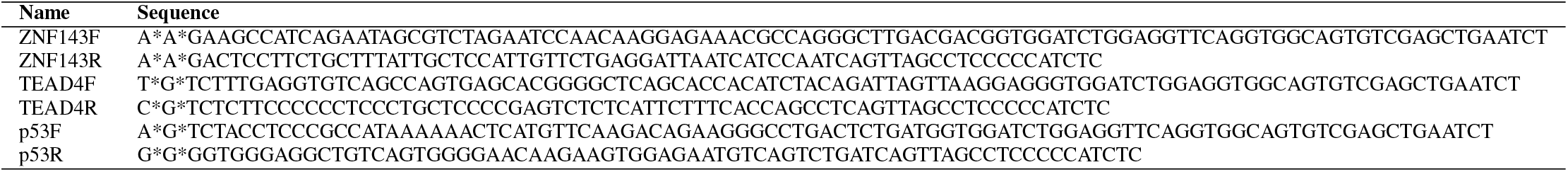
PCR Homology Donor Construct Primers

**Table 2.**
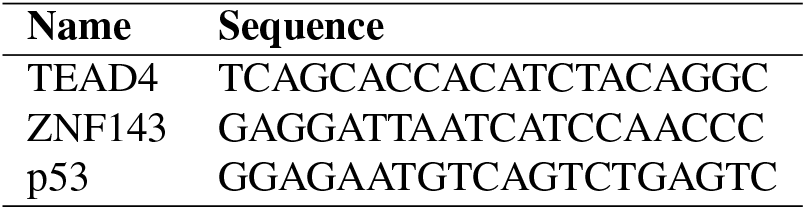
sgRNA Primers (DNA target sequence)

### EGFP-ARF16-PB1 stable cell lines

We transfected plasmids expressing NLS-ARF16-PB1 fused with eGFP at the C-terminus or eGFP-NLS-ARF16-PB1K944A, D994A, D998A mutant into each of the following: HEK293T, HEK293T-TIR1, and HEK293T-TIR1 cells where either TEAD4 or ZNF143 were AID-tagged. Cells were expanded for one week and GFP-sorted iteratively (three times) until we obtained a stable population of GFP expressing cells.

### Immunoprecipitation and immunoblotting

HEK293T or HEK293T-TIR1 cells were co-transfected with mCherry-NLS-AID and eGFP-ARF16-PB1 or eGFP-ARF16-PB1 mutant plasmids. The mutant eGFP-ARF16-PB1 plasmid was co-transfected at a concentration three times higher than the wild type to get comparable expression of the mutant protein. The co-IP data from Figure 4 was generated by separately transfecting mCherry-AID into HEK293T, HEK293T-eGFP-ARF16-PB1, and HEK293T-TIR1-eGFP-ARF16-PB1 cells. Cells were lysed 24 hours post-transfection in a buffer containing 50 mM Tris (pH 7.5), 150 mM NaCl, 0.5% NP-40, 1 mM EDTA, and protease and phosphatase inhibitors for 30 minutes on ice and then sonicated 30 seconds on and 30 seconds off for 15 cycles. Lysate was clarified by centrifugation for 15 minutes at 12000g in 4°C. Anti-GFP antibody conjugated magnetic beads (Chromotek, gtma-10) or anti-mCherry conjugated affinity gel (Biolegend, 689502) were blocked with 1% BSA for 15 minutes. The beads were incubated with clarified lysate for 1.5 hours at 4°C. The immunoprecipitate was recovered with DynaMag racks or by centrifuging at 3000xg for 1 minute. Beads were washed for 5 minutes three times in the lysis buffer and 60 *μ*l 2X Laemmli buffer was added directly to the beads. The complex was heat denatured at 95°C for 5 minutes. We used the following antibodies for the western blots: anti-GFP (gift from Daniel Foltz, Northwestern University), anti-mCherry (1:5000, Rabbit, abcam, ab183628), ZNF143 (1:5000, H00007702-MO1, Abnova), TEAD4 (1:1000, sc-101184, Santa Cruz), p53 (1:1000, DO1, Santa Cruz), anti-TIR1 1: 10,000, gift from Masato Kanemaki, Osaka University), *β*-Actin (1:5000, Sigma A1978), *β*-Tubulin (AA2, gift from Todd Stukenberg, University of Virginia), CENP-I (Rabbit, gift from Todd Stukenberg, University of Virginia). Regression lines for kinetic data were fit using GraphPad Prism (Motulsky and Christopoulos 2004).

### Drug treatment

Cells were treated with 10 *μ*M MG132 for 4.5 hours to test whether AID-tagged proteins were degraded through the proteasome pathway. A stock of 50 mM Auxin was diluted to a final concentration of 500 *μ*M auxin in the culture media. The 50 mM stock was solubilized in DMSO for the 3 hour auxin treatment PRO-seq experiments using the progenitor line. For all other experiments, the 50 mM auxin was solubilized in water. A Degradation rate upon auxin treatment was measured by treating cells with 10 *μ*g/ml cycloheximide and 500 *μ*M Auxin and collecting samples at every 15 minutes for 4 hours.

### PRO-seq library preparation

Cell permeabilization was performed as previously described (Mahat et al. 2016). Cells were collected in 10 ml ice cold PBS after trypsinization and then collected and washed in 5 ml buffer W (10 mM Tris-HCl pH 7.5, KCl 10 mM, Sucrose 150 mM, 5 mM MgCl_2_, 0.5 mM CaCl_2_, 0.5 mM DTT, 0.004 units/ml SUPERaseIN RNase inhibitor (Invitrogen), Protease inhibitors (cOmplete, Roche)). The washed cells were then permeabilized with buffer P (10 mM Tris-HCl pH 7.5, KCl 10 mM, Sucrose 250 mM, 5 mM MgCl_2_, 1 mM EGTA, 0.05% Tween-20, 0.1% NP40, 0.5 mM DTT, 0.004 units/ml SUPERaseIN RNase inhibitor (Invitrogen), Protease inhibitors (cOmplete, Roche)) for 3 minutes. Cells were washed again with 10 ml buffer W before transferring into 1.5 ml tubes using wide bore pipette tips. Finally, cells were re-suspended in 500 *μ*l buffer F (50 mM Tris-HCl pH 8, 5 mM MgCl_2_, 0.1 mM EDTA, 50% Glycerol and 0.5mM DTT). After counting the nuclei, we generated 50 *μ*l aliquots with approximately 3-5 × 10^5^ cells that were snap frozen in liquid nitrogen and stored at −80°C. All centrifugations were done at 500xg for 10 ml conical tubes and 2000xg for 1.5 ml tubes at 4°C and all buffers were maintained on ice. PRO-seq libraries were prepared as described previously (Duarte et al. 2016), with the following modifications. The libraries were amplified by PCR for a total of 10 cycles. We performed 5*′* de-capping using RppH, 5*′* hydroxyl repair, 5*′* adapter ligation, and reverse transcription while the 3*′* RNA biotin moiety was bound to magnetic streptavidin beads. We added an eight base random unique molecular identifier to the 5*′* end of the adapter that is ligated to the 3*′* end of the nascent RNA. We did not perform any size selection because we were willing to tolerate excessive adapter/adapter ligation products to ensure that our libraries were not biased against short nascent RNA insertions.

### PRO-seq analyses

We removed adapters from the paired end 1 or single end reads using `cutadapt` (Martin 2011). Each 3*′* adapter harbored an 8 base unique molecular identifier (UMI). We removed PCR duplicates based on the UMIs using `fqdedup` (Martins and Guertin 2018). We trimmed UMIs with `fastx_trimmer` (Gordon 2010) and we implemented `fastx_reverse_complement` to generate the reverse complement sequence (Gordon 2010). We aligned reads to *hg38* with `bowtie2` (Langmead et al. 2009). We sorted aligned *BAM* files using `samtools` (Li et al. 2009). We used `seqOutBias` to generate *bigWig* files (Martins et al. 2017). We used the `bigWig` R package (Martins 2014) and UCSC Genome Browser Utilities (Kent et al. 2010) to query *bigWig* files within genomic coordinates. `Bedtools` was used to parse genomic coordinate files and query for overlapping regions (Quinlan and Hall 2010). Differential nascent transcript abundance was measured by `DESeq2` (Love et al. 2014). Bidirectional transcription was identified using `dREG` (Wang et al. 2019). `MEME` was used for *de novo* motif discovery within dREG-identified regulatory elements that change upon ZNF143 depletion; `TOMTOM` matched the ZNF143 motif (Bailey et al. 2015b) to a database that is curated by `HOMER` (Heinz et al. 2010). All the analysis details and code are available at https://github.com/mjg54/znf143_pro_seq_analysis. Raw sequencing files and processed *bigWig* files are available from GEO accession record GSE126919.

### Model formulation

The dynamics for the concentrations or densities of RNA Polymerases at pausing regions and gene bodies, defined as *p* and *b*, are described as follows:

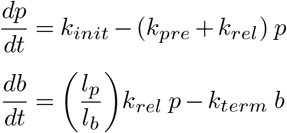

where *k*_*init*_ is the rate of transcription initiation, *k*_*pre*_ is the rate constant for the release of a paused Pol II as transcription proceeds into productive elongation in the gene body, and *k*_*term*_ is the rate constant for transcription termination. The term *l*_*p*_/*l*_*b*_ is a ratio of the relative DNA segment lengths that is applied to adjust the gene body concentration based on the larger amount of DNA in the gene body as compared to the pause region. The term *k*_*init*_ implicitly accounts for the product of the unbound Pol II concentration and the rate constant for initiation. Concentration is considered as number of RNA polymerases per length of DNA. Thus this concentration is referred to as a density. We consider this model as though the units are dimensionless for the analysis of how specific rates influence the relative Pol II quantities at pause sites and within the gene bodies. The steady state levels (*p*_*ss*_, *b*_*ss*_) are found by setting *dp/dt* = *db/dt* = 0. Because understanding the effects of the relative pause region length is not our focus, we set *r* = *l*_*p*_/*l*_*b*_ for the parameter sensitivity analyses.

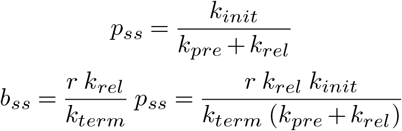

The pause index is the relative density of reads in the pause region compared to the gene body (*P*_*i*_ = *p*_*ss*_/*b*_*ss*_). The pause index is only dependent on *k*_*term*_ and *k*_*rel*_, the rate constants for the termination of transcription and release into productive gene body elongation:

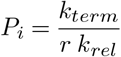

### Parameter sensitivity analysis

To examine the effects of each parameter on the pause and body concentrations, we considered high and low parameter values (*k*^(*hi*)^ and *k*^(*lo*)^) and computed the changes in *p*_*ss*_ and *b*_*ss*_. The following notation documents the change in *p*_*ss*_ when *k* is changed from a relatively high to a relatively low value: ∆*p*(*k*) = *p*_*ss*_(*k*^(*hi*)^) − *p*_*ss*_(*k*^(*lo*)^)). First, we evaluate the effects of the transcription initiation rate (*k*_*init*_) on the steady state pause region and gene body Pol II concentrations.

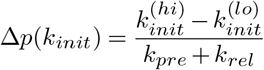

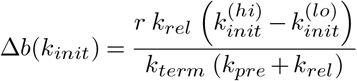

Note that *k*^(*hi*)^ *> k*^(*lo*)^ by definition, so that ∆*p*(*k*_*init*_) *>* 0 and ∆*b*(*k*_*init*_) *>* 0. Therefore, increasing the rate of transcription initiation will result in both pause region and gene body increases. We present results of single parameter changes that can also be considered using a standard sensitivity analysis. For this example the sensitivities are computed as follows:

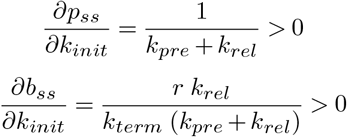

In general, ∆*p*(*k*) ~ (*∂p/∂k*)(*k*^(*hi*)^ – *k*^(*lo*)^) for small changes in *k* and *sign*(∆*p*(*k*)) = *sign*(*∂p/∂k*). Next we consider the effects of varying the rate constant for premature pause release (*k*_*pre*_).

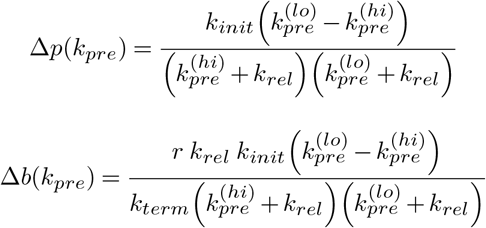

These results show that increasing the rate constant for premature pause release will decrease both pause region and gene body Pol II concentrations (∆*p*(*k*_*pre*_) < 0 and ∆*b*(*k*_*pre*_) < 0). We next demonstrate that an increase in the rate constant for the release of Pol II into the gene body (*k*_*rel*_) decreases *p*_*ss*_ and increases *b*_*ss*_:

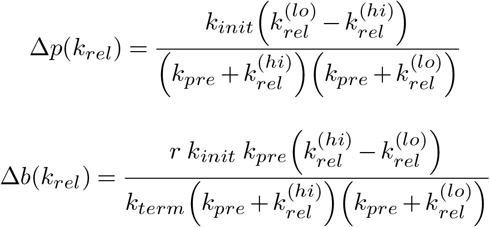

It is interesting to note that if there is no premature pause release (i.e., *k*_*pre*_ = 0), the model predicts that a change in the rate of pause release into transcriptional elongation will not affect the concentration of RNA polymerases in the gene body, if all other factors are identical (i.e., ∆*b*(*k*_*rel*_) = 0 for *k*_*pre*_ = 0). Finally, we evaluate the effect of modifying the rate constant for the termination of transcription:

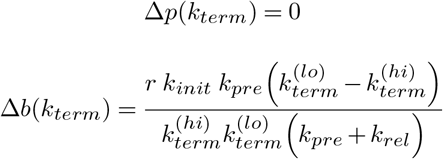

These results shows that an increase in the rate constant for transcriptional termination does not affect the steady state level of paused Pol II but will decrease the gene body concentration of RNA polymerases (∆*p*(*k*_*term*_) = 0, ∆*b*(*k*_*term*_) < 0). The results from our parameter sensitivity analyses are illustrated in Table 3 where the entries for *p*_*ss*_ and *b*_*ss*_ indicate the effect of increasing each parameter.

**Table 3.**
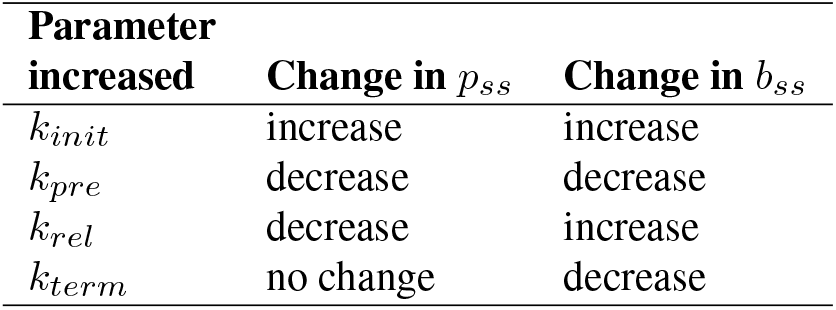
Parameter sensitivity summary

### Relative effects of transcriptional initiation and premature release on Pol II distribution

Our preceding analyses show that consistent changes in the pause region and gene body (i.e., both either increase or decrease) are only observed following changes in the rate of transcriptional initiation or the rate constant for premature pause release (*k*_*init*_, *k*_*rel*_). Our experimental data show that repressed genes show decreases in both the pause region and the gene body region. These decreases in Pol II density were greater in the pause region in comparison to the gene body. Next, we document the conditions under which the effects of varying the rates of initiation and premature release of the paused Pol II are greater for the pause region as compared to the gene body (e.g., ∆*p*(*k*_*init*_) > ∆*b*(*k*_*init*_)). Changes in transcriptional initiation are set as 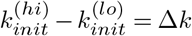:

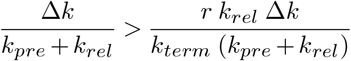

This condition is satisfied for 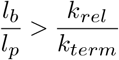 (recall *r* = *l*_*p*_/*l*_*b*_). In general, *l*_*b*_/*l*_*p*_ is a large value because the pause region (*<*100 bp) is much smaller than the gene body (approximately > 10kb. So *k*_*rel*_/*k*_*term*_ < *l*_*b*_/*l*_*p*_ ~ 100 must be obtained. For changes in premature release, the condition is as follows:

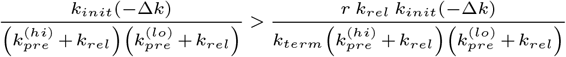

This relation leads to the same constraint observed for *k*_*init*_: *k*_*rel*_/*k*_*term*_ < *l*_*b*_/*l*_*p*_. Recall that the pause index is defined as *P*_*i*_ = *k*_*term*_/*r k*_*rel*_ = (*l*_*b*_/*l*_*p*_)(*k*_*term*_/*k*_*rel*_). Therefore, *P*_*i*_ > 1 for the same condition that constrains ∆*p*(*k*_*pre*_) > ∆*b*(*k*_*pre*_) for changes in *k*_*init*_ and *k*_*pre*_: *k*_*rel*_/*k*_*term*_ < *l*_*b*_/*l*_*p*_. This demonstrates that, for an arbitrary change in either the initiation rate or the premature pause release rate constant, the effect of the change in the pause region will be greater than the change in the gene body region whenever the pause region Pol II concentration is greater than that at the gene body. Further, the effect ratio is identical for changes in *k*_*init*_ and *k*_*rel*_ and is equal to the value of the pause index: ∆*p*/∆*b* = *P*_*i*_.

### Pause region and gene body model visualization

We aimed to generate plots in which steady state levels of pause region and gene body concentration were imposed upon the peak and flat regions of a profile that is characterized by an exponential approach towards the peak followed by an exponential decay towards a stable plateau. We used a sum of exponential functions for the waveform:

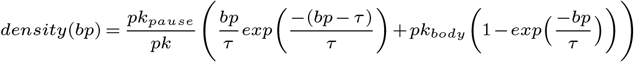

where *bp* (base pairs) is the independent variable and *τ* is the exponential decay constant. The parameter *pk* is set to the root of the derivative as shown below so that *pk*_*pause*_ determines the peak of the waveform. The parameter *pk*_*body*_ is set such that the asymptotic gene body region decays to a desired level. The derivative of this waveform is

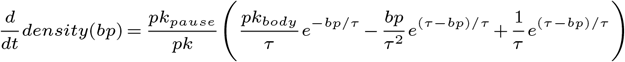

and the root of the derivative is given by the value of *bp* at the peak of the waveform (*max*(*density*)):

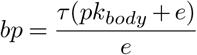

For *bp* >> *τ*, the gene body level assymptotically approaches *pk*_*body*_ *pk*_*pause*_/*pk*. We implicitly determine a value for *pk*_*body*_ that will give a gene body level of a desired level. We select a value of *pk*_*body*_ that will give a plateau *pk*_*body*_ *pk*_*pause*_/*pk* of choice for a given setting of the pause peak *pk*_*pause*_. After implicitly finding a value for *pk*_*body*_ that produces the gene body level of choice, the waveform is produced.

## ACKNOWLEDGEMENTS

This work was funded by R35-GM128635 to MJG and R21-HG009021 to LC. We thank Dr. Ricardo Henriques for making the bioR*χ*iv L^A^T_E_X template publicly available. We thank Arun Dutta and Drs. Anindya Dutta, Todd Stukenberg and Prasad Trivedi for discussion and comments. Dr. Ryan Lewis from http://scidelight.com/ illustrated Figures 1A, 8, S1, and S3D.

## AUTHOR CONTRIBUTIONS

KMS conceived of the ARF-AID hypothesis. KMS and BDM performed the experiments. WA implemented the mathematical models. FMD, LJC, and MJG developed the PRO-seq protocol. KMS and MJG conceptualized and developed the project. KMS and MJG designed the experiments. MJG and KMS analyzed the data. MJG supervised the project. MJG acquired the resources and funding. KMS, WA, and MJG wrote the original draft. KMS, BDM, WA, FMD, LJC, and MJG edited the manuscript.

---

**Fig. S1.**
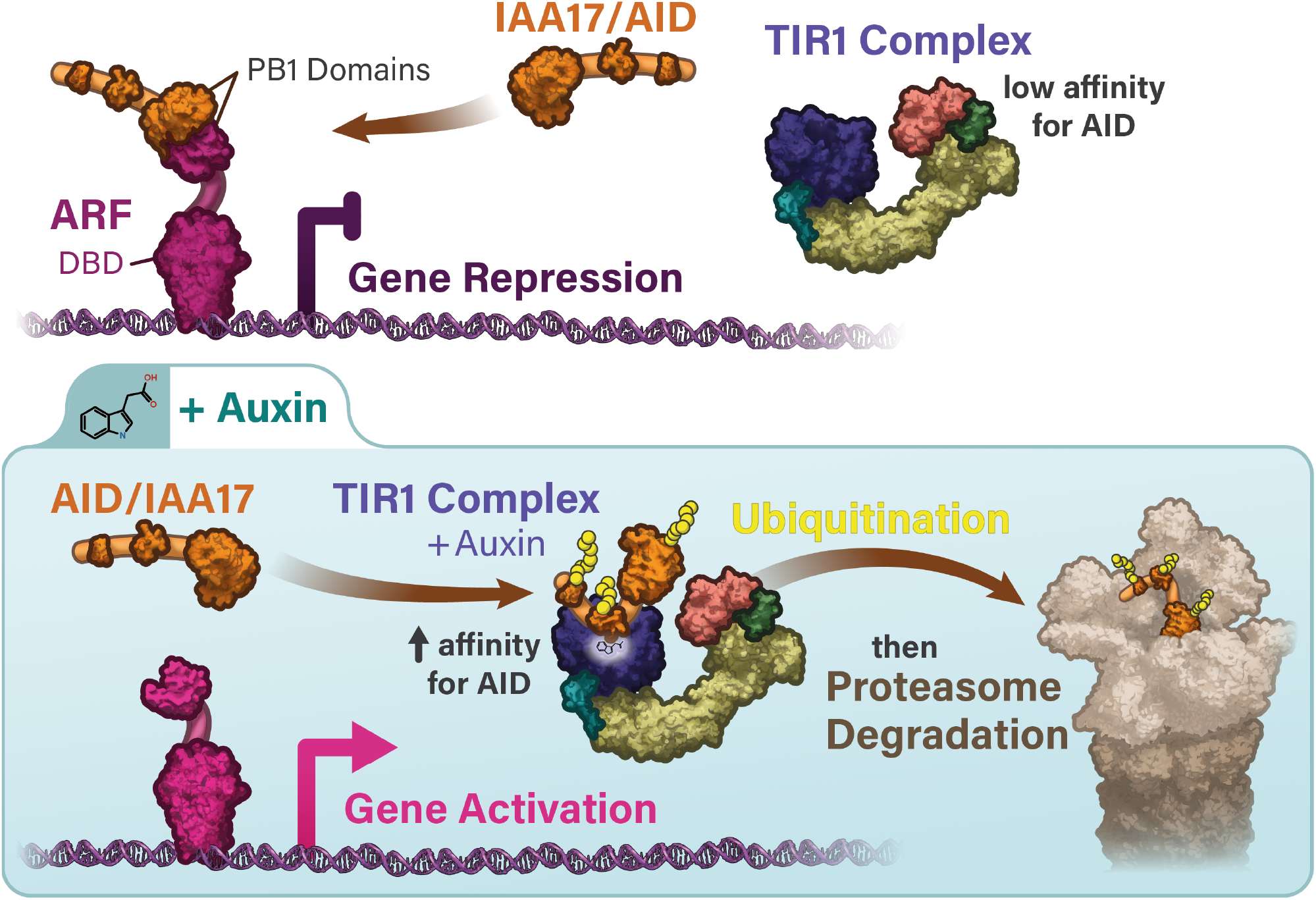
ARF is critical for the proper regulation of the auxin-induced transcriptional response in plants. In the absence of auxin, IAA binds the PB1 domain of ARF. Auxin binds to TIR1 and drives strong association of TIR1 with IAA to mediate ubiquitination and degradation.

**Fig. S2.**
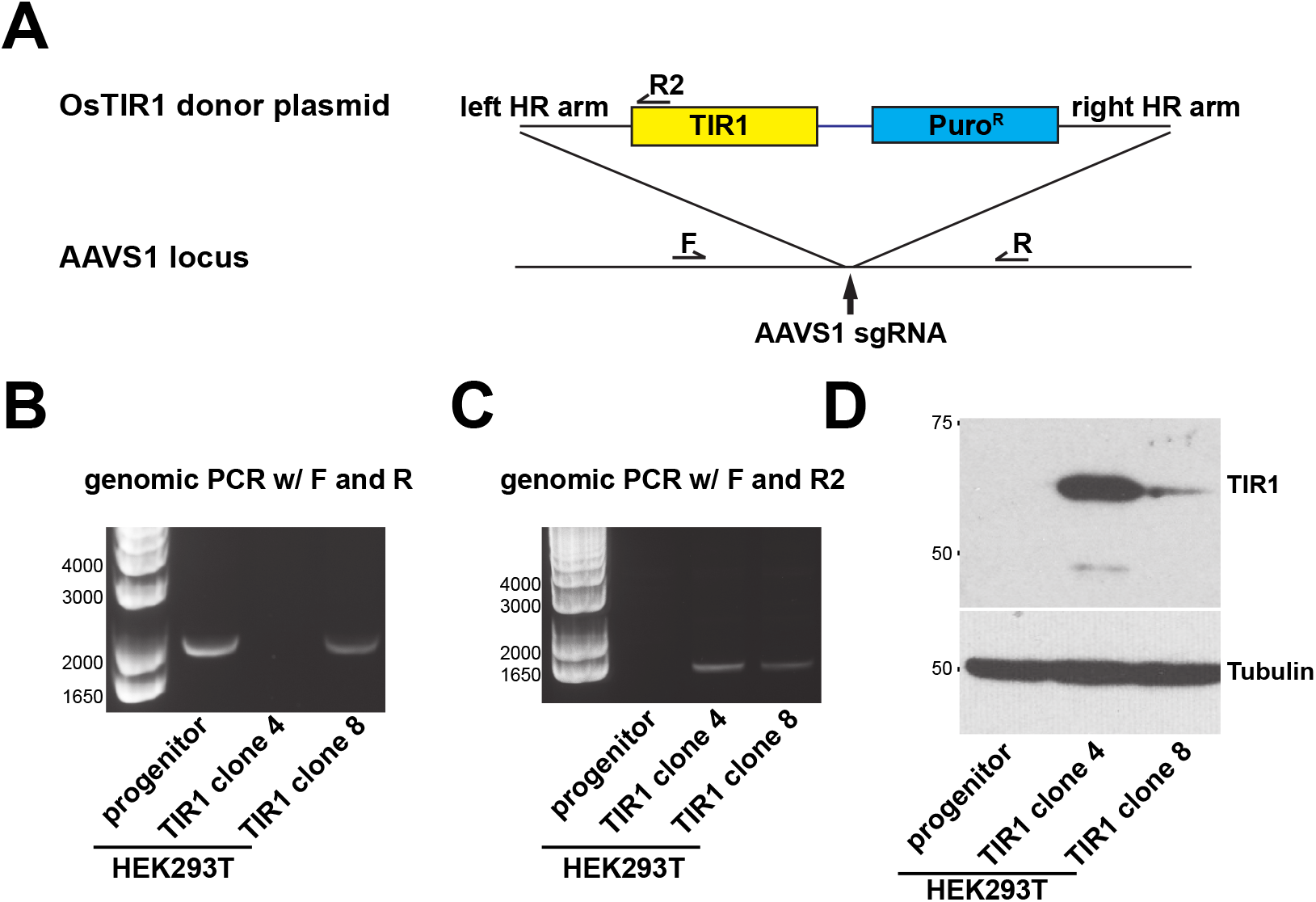
TIR1 was integrated into the AAVS1 locus of HEK293T cells. A) A schematic of the strategy to integrate CMV-driven TIR1 into the AAVS1 locus of the HEK293T cells using the CRISPR-Cas9 system. Primers F and R2 generate an amplicon if the construct is inserted. F and R primers flank the insert and will amplify if the construct does not insert into a copy of AAVS1. B) At least one copy of AAVS1 contains the insertion in both clone 4 and clone 8. C) All copies of AAVS1 contain the insertion in clone 4, but clone 8 is heterozygous. D) Western blotting indicates that the TIR1 protein is expressed in both clones.

**Fig. S3.**
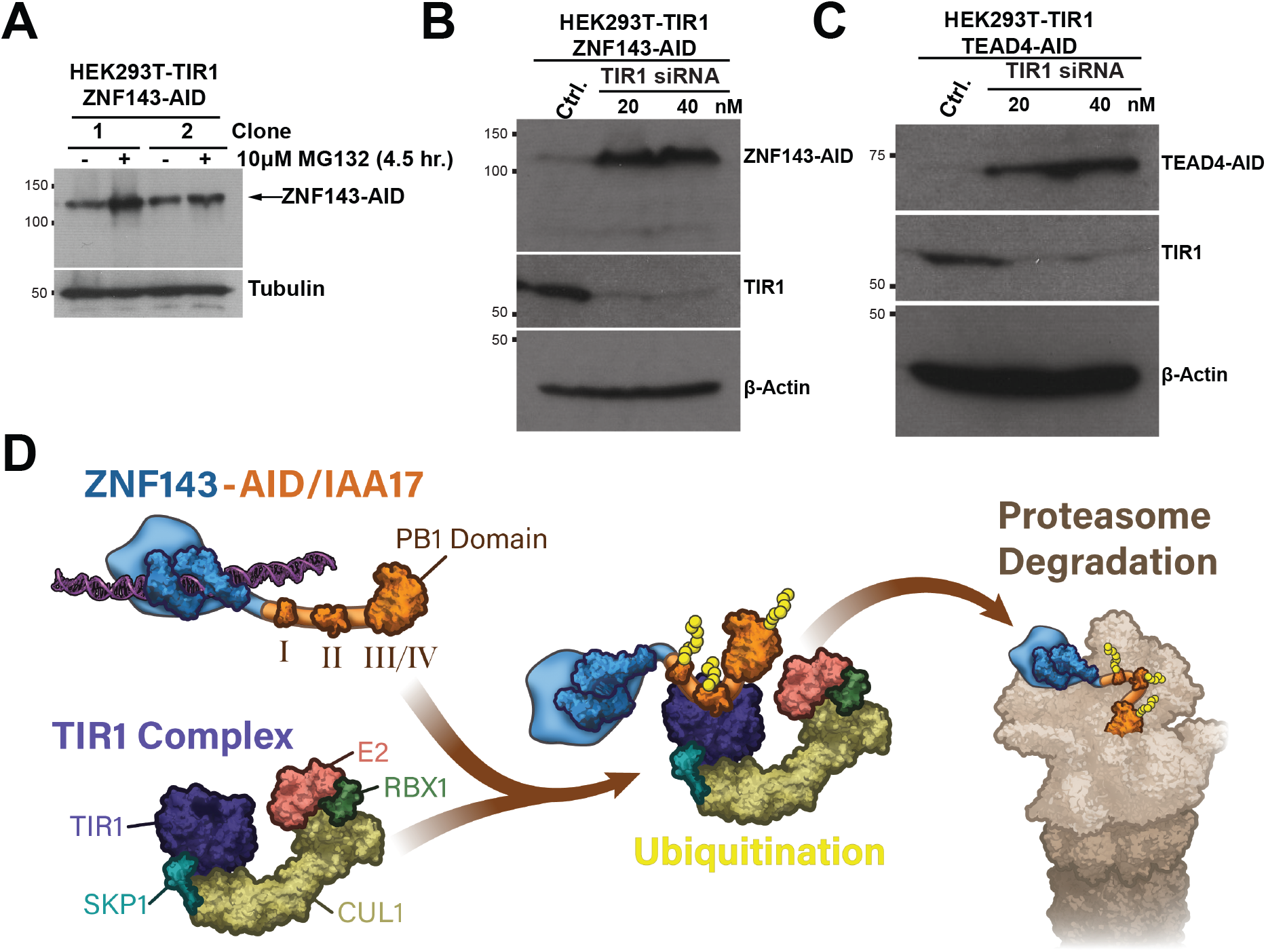
Chronic auxin-independent depletion is proteasome dependent. A) Two independent clones (1 and 2) of ZNF143-AID tagged cells were treated with MG132 for four and a half hours. ZNF143 protein levels increase after MG132 treatment. B&C) TIR1 depletion by siRNA knockdown stabilizes ZNF143-AID and TEAD4-AID. D) An auxin-independent interaction between ZNF143-AID and TIR1 is proposed to drive chronic proteasomal degradation.

**Fig. S4.**
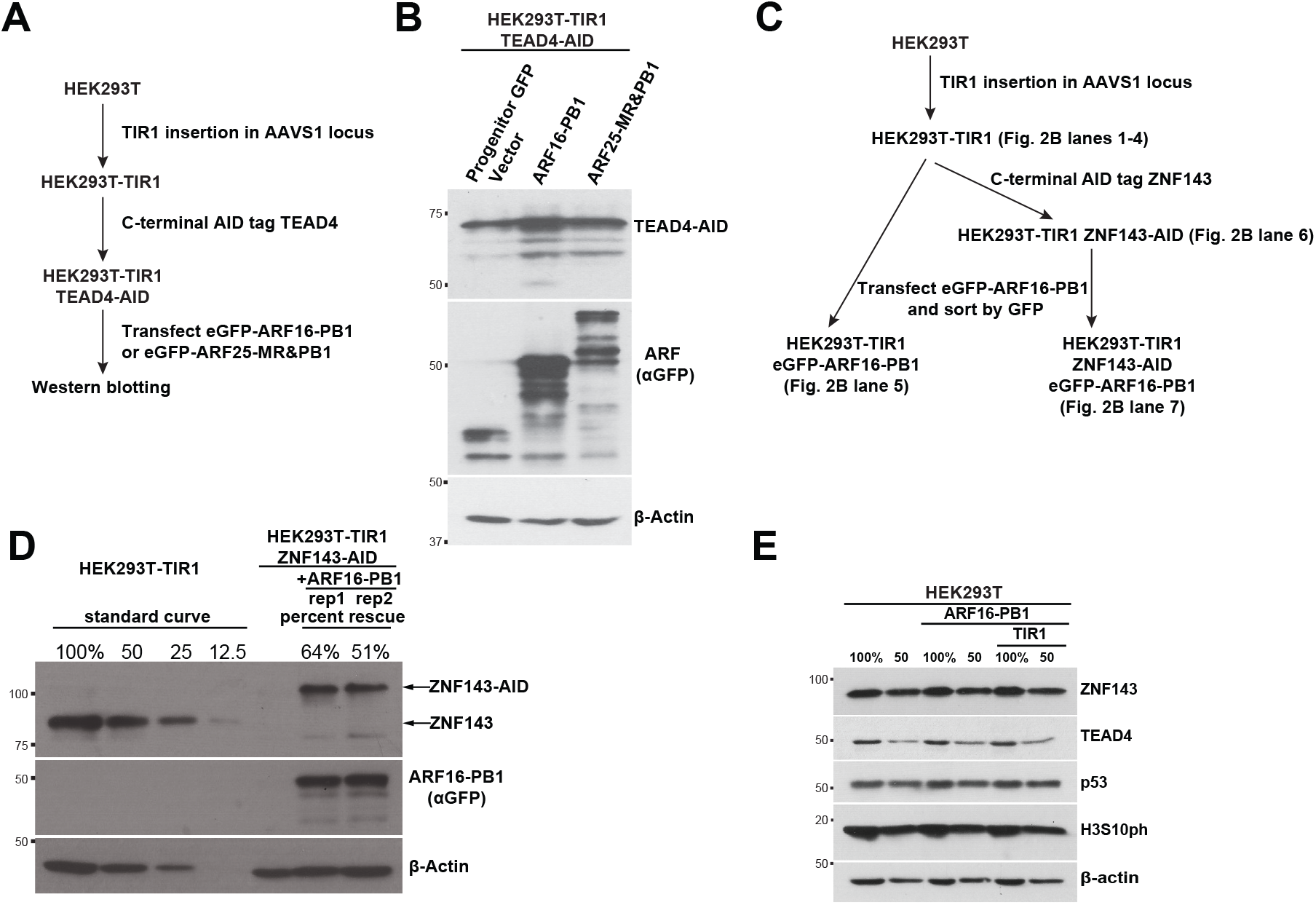
Expression of the ARF domains rescues chronic protein depletion, but does not detectably affect protein levels in the absence of AID-tagging. A) A schematic of the strategy to express ARF (PB1 domain of ARF16 or the MR and PB1 domain of ARF25) within HEK293T cells that express TIR1 and AID-tagged TEAD4. B) TEAD4-AID protein is stabilized after expression GFP-ARF16-PB1 and GFP-ARF25-MR-PB1, but not stabilized by the empty vector that expresses GFP. C) The illustrated workflow highlights the source of each lane in Figure 2B. D) Expression of the ARF16 PB1 domain rescues ZNF143 levels to 51% to 64% the level of untagged ZNF143. E) ARF16-PB1 and TIR1 expression had no effect on the expression of ZNF143, TEAD4, and p53; actin and phosphorylated H3S10 are loading controls.

**Fig. S5.**
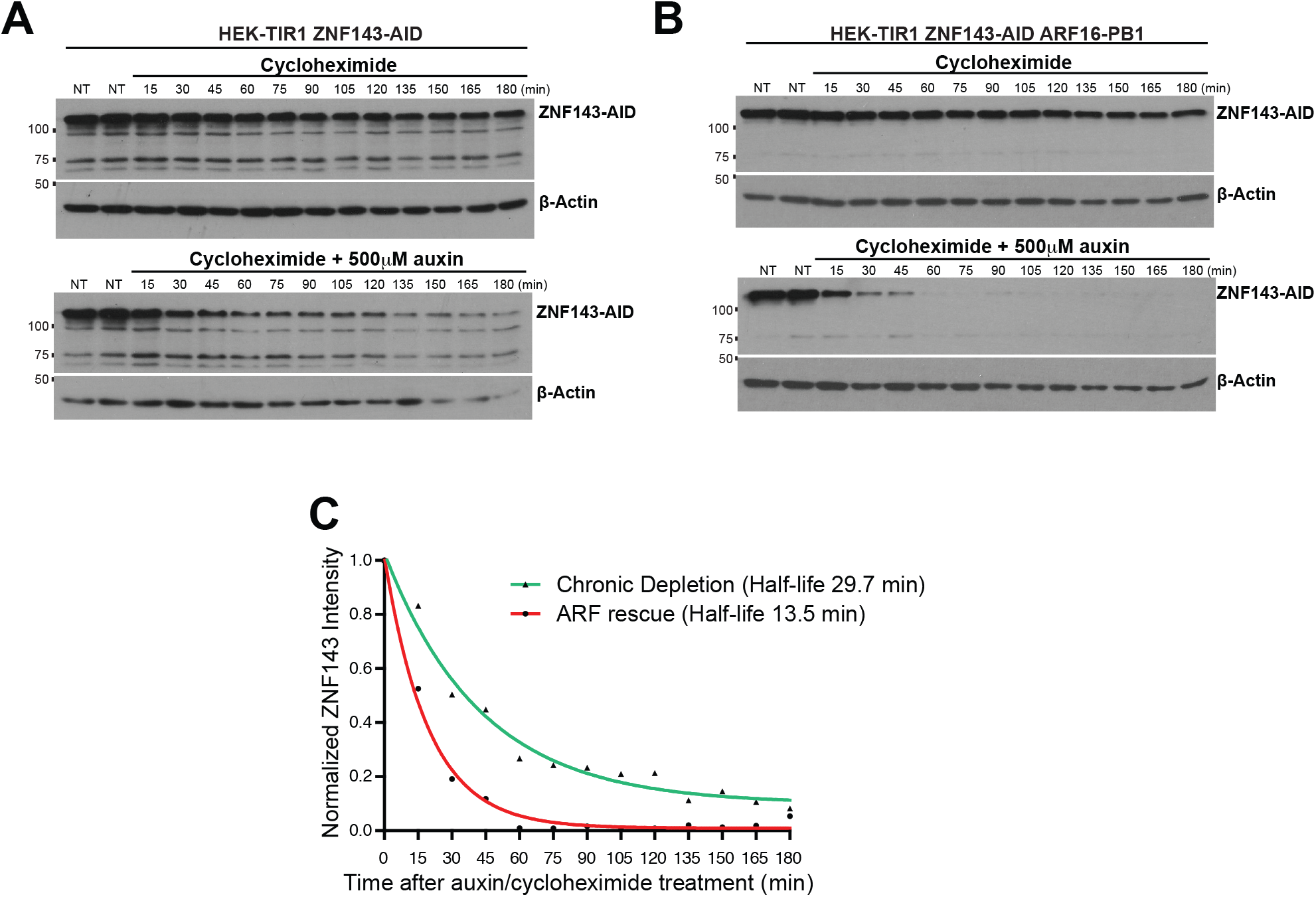
ARF rescue increases the rate of auxin dependent degradation of AID tagged protein. HEK293T-TIR1 ZNF143-AID cells (A) and ARF-16-rescued cells (B) were treated with cycloheximide alone, or cycloheximide in combination with auxin, to compare degradation kinetics upon inhibiting translation. C) Densitometric quantification measurements of ZNF143-AID bands from the lower blots in panel A (green trace) and panel B (red trace) were plotted and fit to a one phase decay equation.

**Fig. S6.**
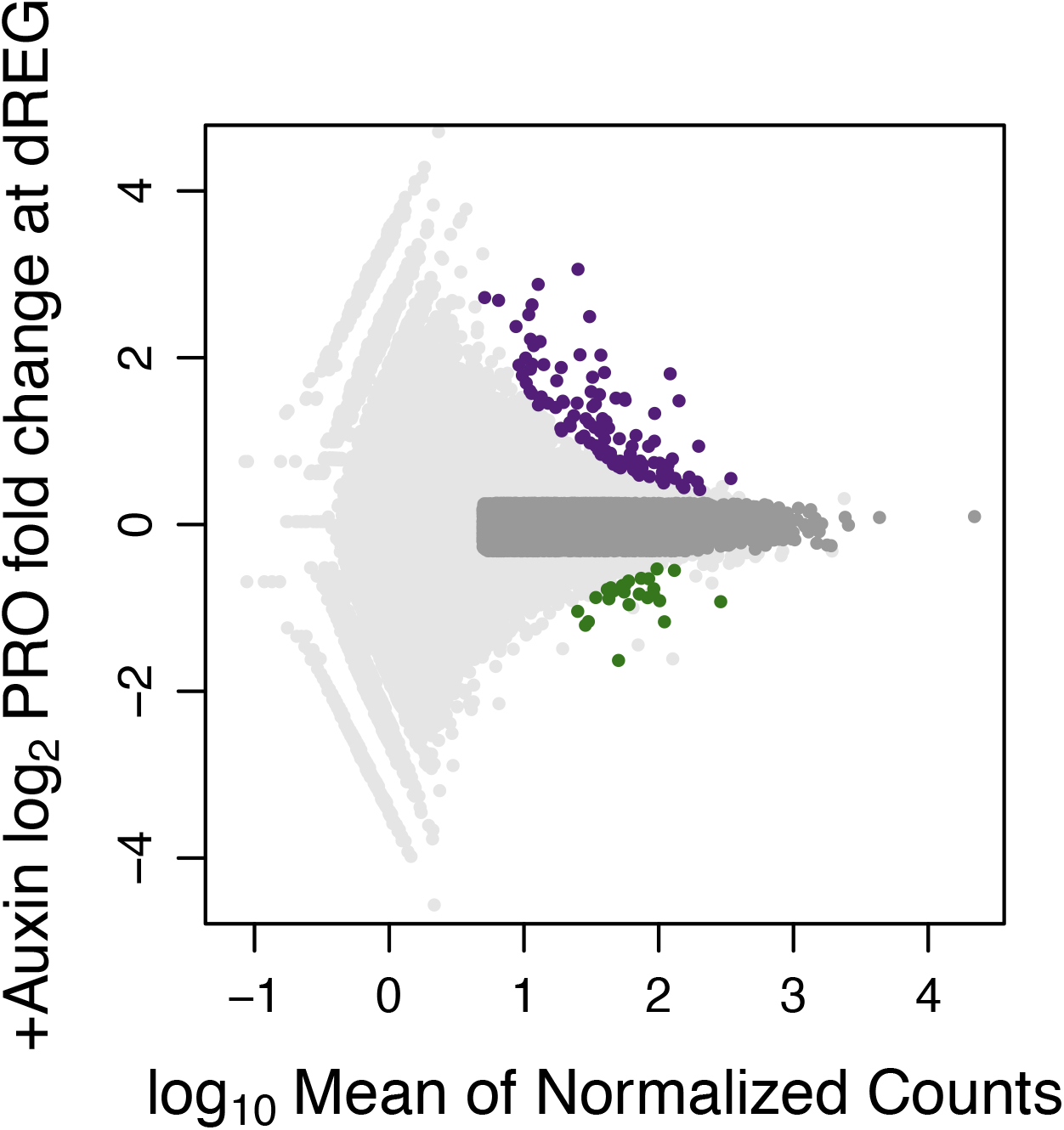
Bidirectional transcription changes modestly at a subset of dREG-defined (Wang et al. 2019) regulatory elements upon ZNF143 depletion.

**Fig. S7.**
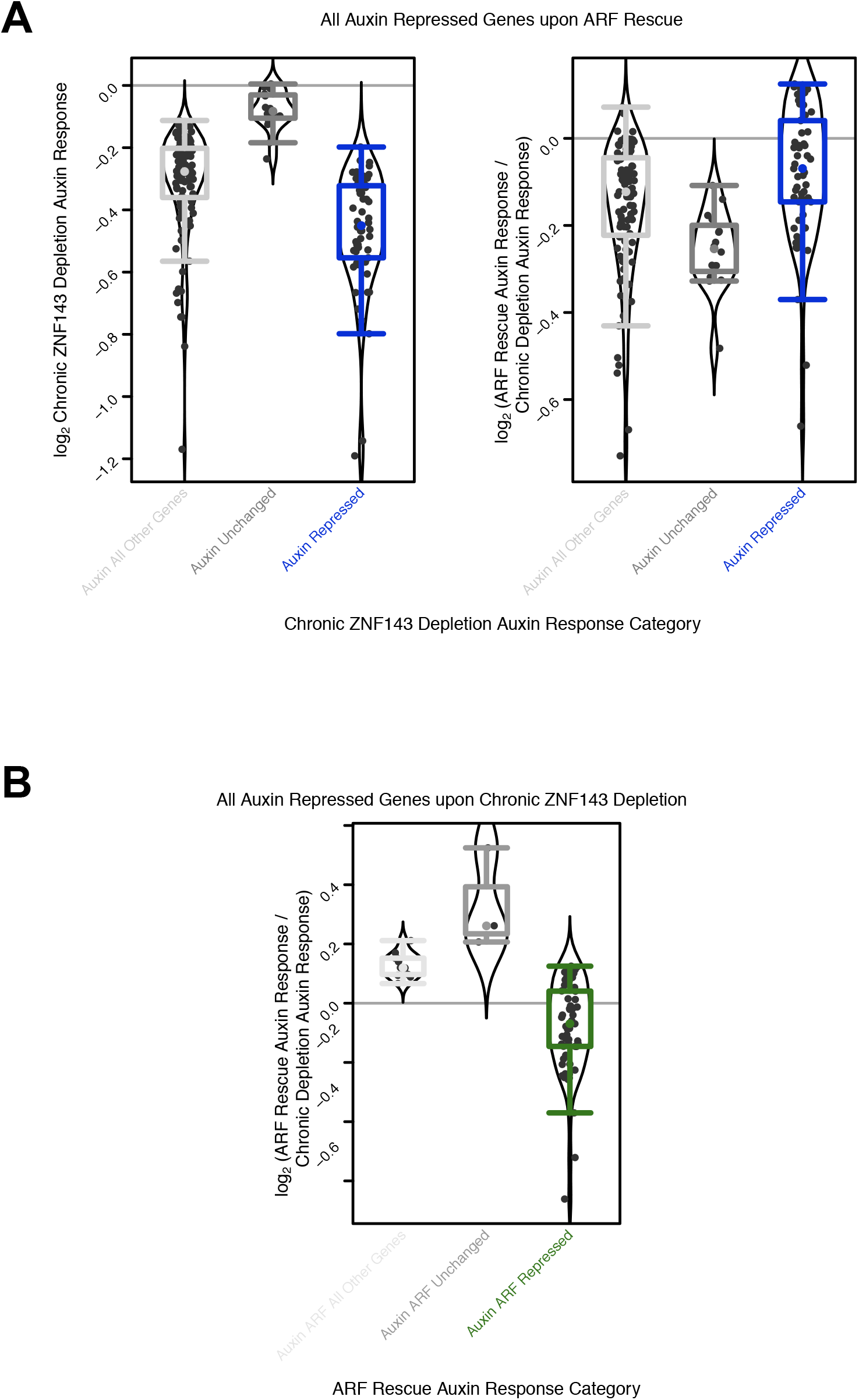
ZNF143-repressed genes decrease expression in the chronic depletion background, but below our sensitivity of detection. A) Despite falling in *unchanged* and *all other genes* classes, the net response upon auxin treatment in the chronic depletion background is consistent in direction with auxin response in the rescue. Genes have a greater magnitude of response in the rescue compared to chronic depletion. B) Repressed genes in the chronic depletion background are not consistently auxin-repressed and we do not consistently observe a greater magnitude of auxin-response in the ARF-rescue. These data suggest that the changes in gene expression in the chronic depletion background may not be ZNF143-specific.

**Fig. S8.**
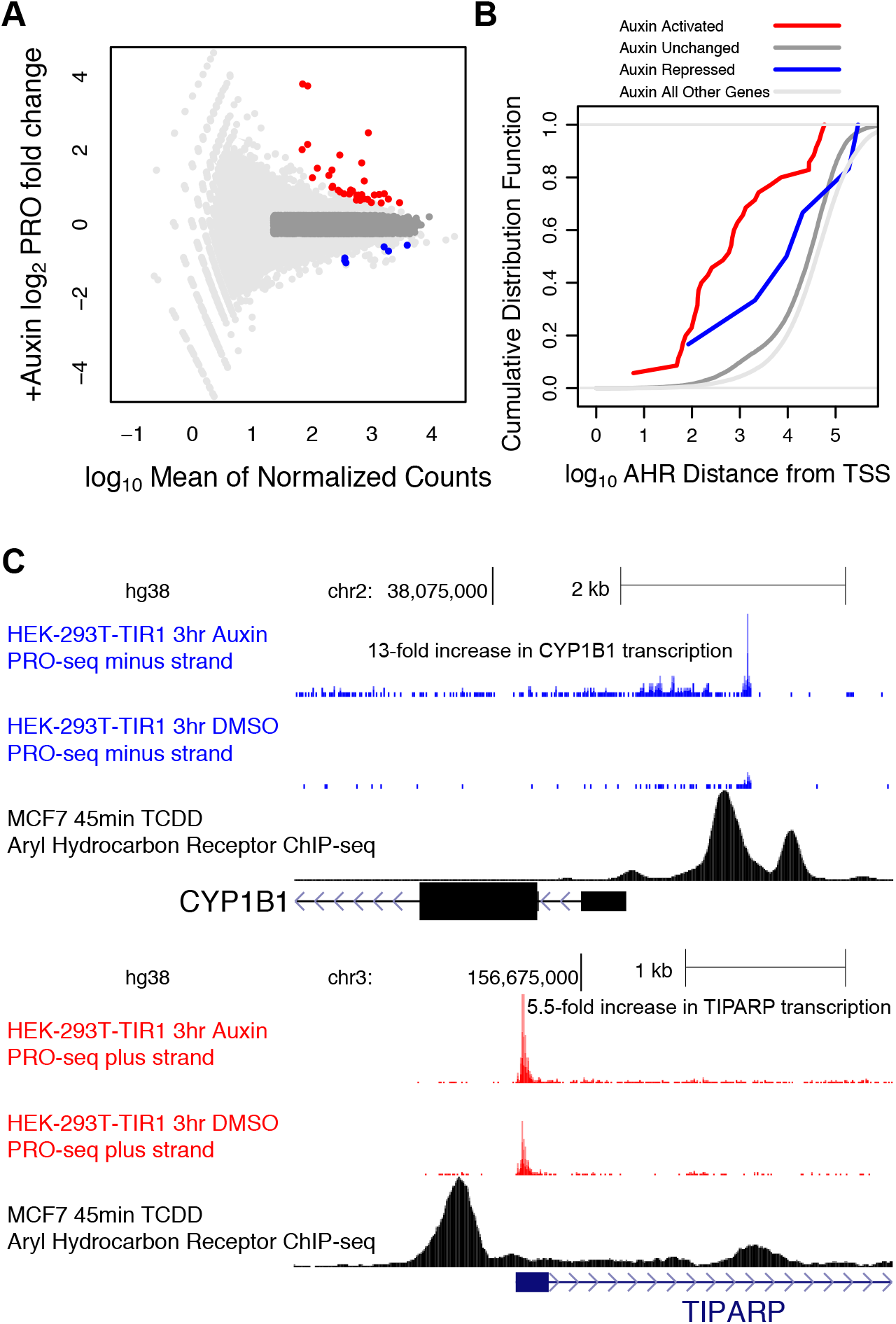
Auxin treatment activates Aryl Hydrocarbon Receptor (AHR) gene targets. A) Auxin treatment activates many genes compared to a DMSO control treatment. B) Auxin-activated genes are, on average, closer to AHR binding sites. C) The genes TIPARP and CYP1B1 are examples of canonical hydrocarbon-responsive genes, both of which are activated by auxin and bound by AHR. Note that the transcription start site for CYP1B1 is upstream of the gene annotation in HEK293T cells.

**Fig. S9.**
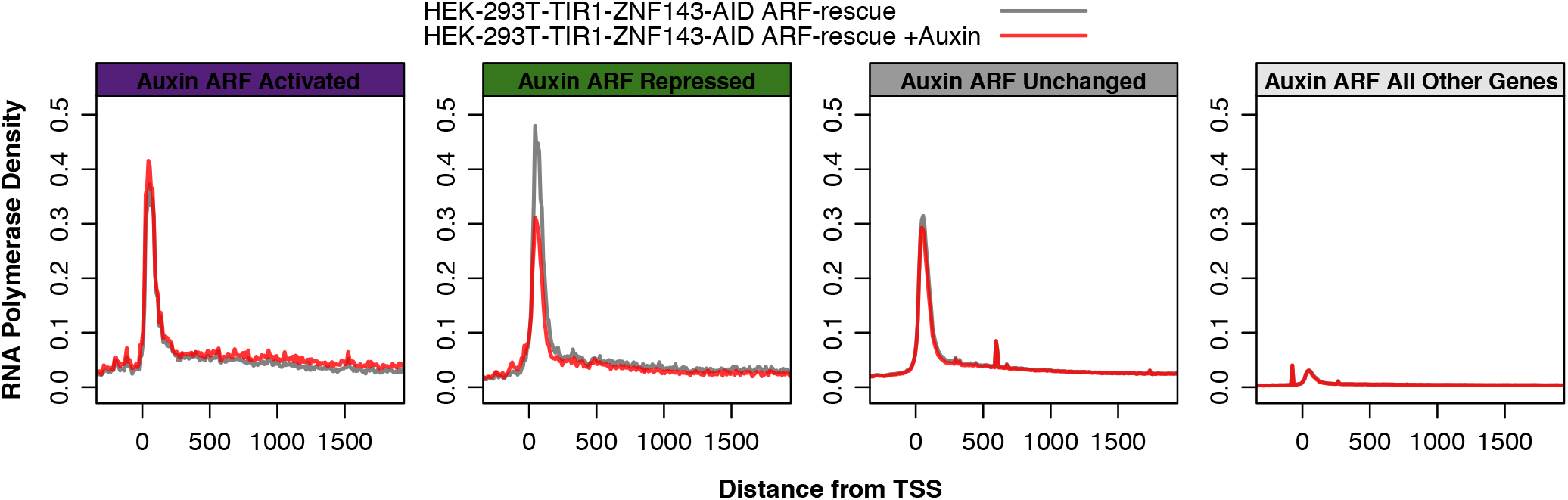
Average RNA Polymerase occupancy plots, or composite profiles, suggest that RNA Polymerase pause density is reduced in the repressed gene class with only modest differences in the other classes.

**Fig. S10.**
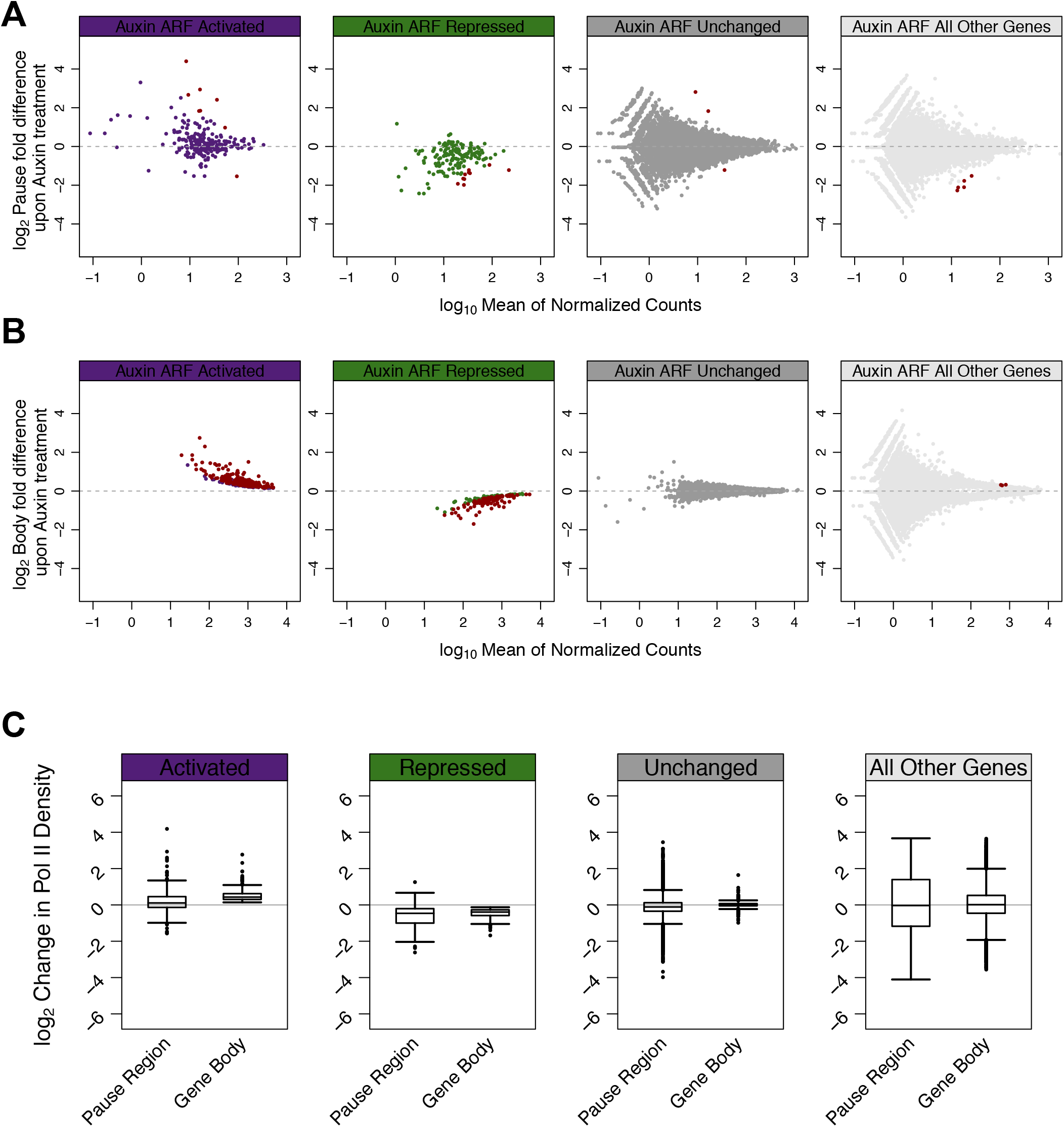
Individual gene analysis indicates RNA Polymerase pause density is reduced in the pause and gene body. A) Pol II pausing is consistently decreased in the auxin-repressed class. Red genes fall below an FDR threshold of 0.01. B) Each panel’s gene class was previously defined by the changes that occur over the entire gene length, which is dominated by the gene body; therefore, red genes (FDR threshold of 0.0001) are consistent with their classification. Note that the x-axis for panels A and B are not identical and only a few genes have a log_10_ mean count above 2 in the pause region for the repressed class, compared to only a few genes below a log_10_ mean count of 2 in the gene body. C) Box and whisker plots collapse the data from panels A&B above.

**Fig. S11.**
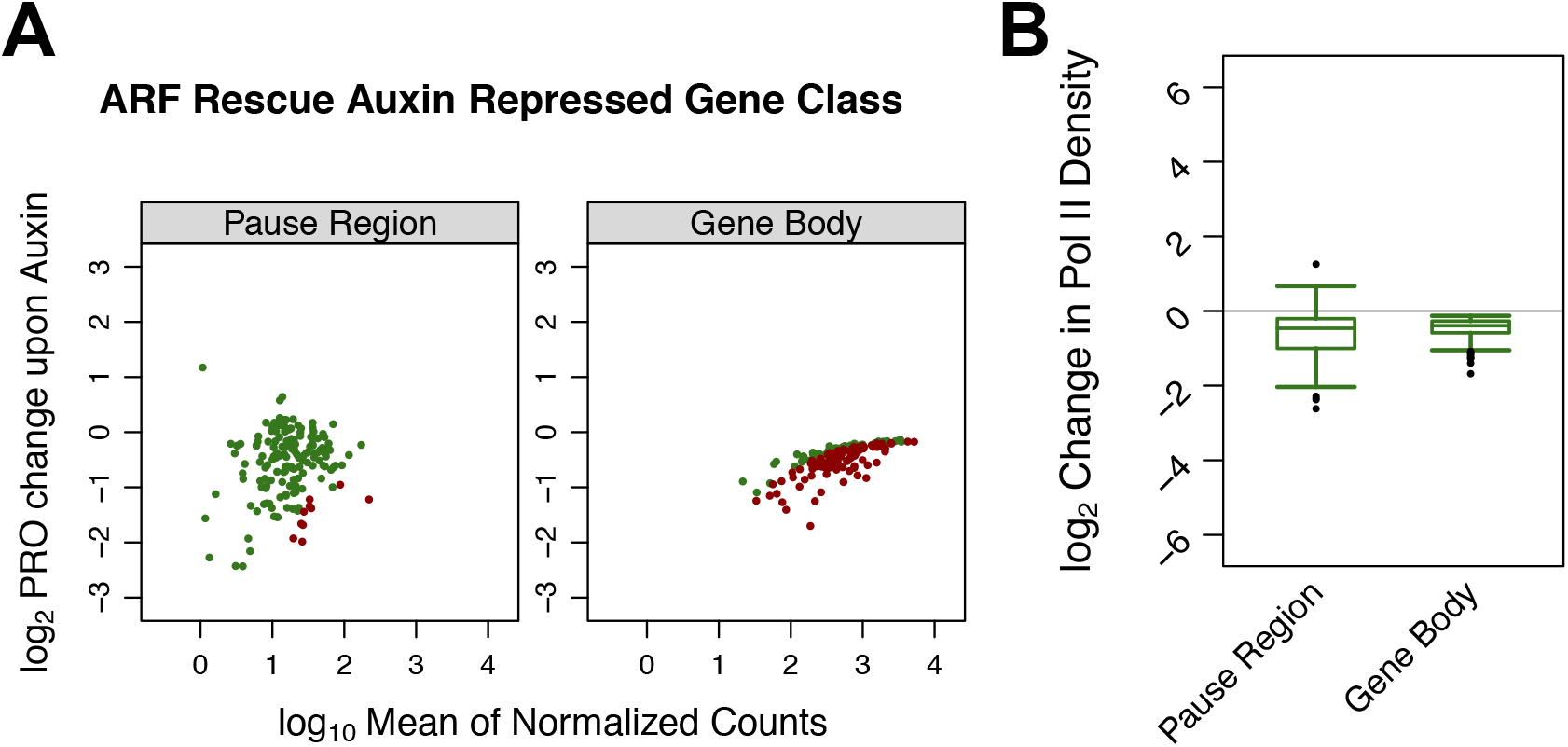
RNA Polymerase pause density is reduced in the pause region and gene body at genes that are repressed upon ZNF143 depletion. The gene body and pause region data for the repressed class from Figure S10 are shown side-by-side.

**Fig. S12.**
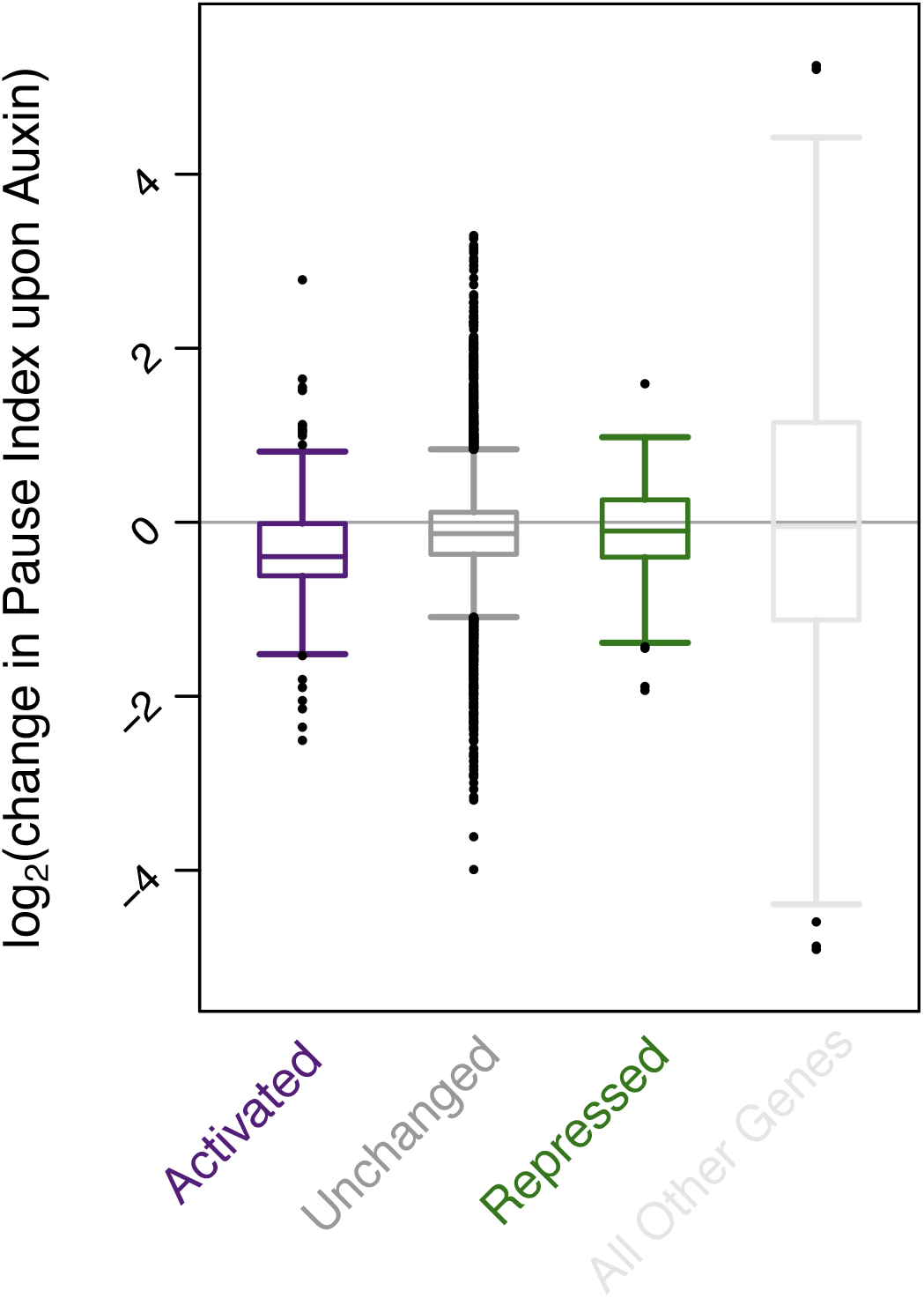
The pause index changes only modestly (log_2_ values of approximately 0) for repressed genes. A modest average decrease in pause index is also observed for the control (unchanged) gene class.

